# Amyloid fibrils in Alzheimer’s disease differently modulate sleep and cortical oscillations in mice depending on the type of amyloid

**DOI:** 10.1101/2025.04.09.647928

**Authors:** Tomomi Sanagi, Masaki Okumura, Yuxi Lin, Shingo Kanemura, Eunyoung Moon, Yunseok Heo, Keiko Takahara, Young-Ho Lee, Tomomi Tsunematsu

## Abstract

Alzheimer’s disease (AD) is characterized by aggregation and deposition of the amyloid-beta (Aβ) protein in patient brains, with aging playing a crucial role through oxidative stress and neuroinflammation. Sleep disturbances are common in patients with AD, and contribute to their cognitive impairment. However, the association between the aggregation of specific Aβ species in particular brain regions and its effect on sleep impairment remains unclear. Here, we investigated the effect of Aβ1–40 (Aβ40) and Aβ1–42 (Aβ42) amyloid fibrils on sleep/wakefulness and cortical oscillations in 3-month old wild-type mice. Aβ42 aggregated faster and demonstrated distinct structural properties compared with Aβ40. Bilateral injections into the dentate gyrus of the hippocampus showed that Aβ42 amyloid fibrils significantly disrupted sleep and cortical activity as well as caused neuronal death, whereas Aβ40 amyloid fibrils mainly affected cortical oscillations and caused minimal neuronal death. These findings shed light on AD-associated sleep disorders, which are differentially affected by the distinct properties of Aβ40 and Aβ42 aggregates.

## Introduction

Alzheimer’s disease (AD) is the most prevalent neurodegenerative disorder, characterized by substantial impairments in behavior and cognition. A neuropathological hallmark of AD is the aggregation of amyloid-beta (Aβ) protein monomers into amyloid fibrils, which subsequently form senile plaques in the extracellular spaces of the brain. Aβ is produced by sequential cleavage of the amyloid precursor protein (APP) by β- and γ-secretases, generating peptides of varying lengths. Aβ1–40 (Aβ40) and Aβ1–42 (Aβ42) are the two major forms of Aβ molecules, although small amounts of other Aβ species, including Aβ1-38, Aβ1-39, Aβ1-41, and Aβ1-43 (Aβ43) have also been reported^1^. The aggregation kinetics and pathways of Aβ is highly sensitive to environmental factors such as ionic strength, lipid membranes, and metal ions^2,3^. In addition to fibrils, Aβ can also form oligomers, which may be act as intermediates along the fibrillization pathway or exist as distinct off-pathway assemblies^4^. While extracellular formation of Aβ plaque is well established, the intraneuronal aggregation of endogenous Aβ or Aβ taken up into cells has been also suggested^5,6^.

Aging, one of the greatest risk factors of AD, plays a crucial role in the aggregation and deposition of Aβ through various molecular and cellular mechanisms. Aging increases the activity of enzymes involved in the processing of APP, and reduces the efficiency of Aβ clearance, thereby leading to the formation and accumulation of Aβ aggregates both inside and outside of cells^7–10^. Moreover, aging increases oxidative stress, which impairs mitochondrial function and weakens antioxidant defense mechanisms, and indirectly promotes Aβ aggregation and accumulation^11–14^. Neuroinflammation induced by aging also indirectly boosts aggregation by stimulating Aβ production through the activation of β- and γ-secretases^15,16^. Furthermore, aging-induced oxidative stress directly leads to Aβ oxidation^14,17–19^. Oxidative stress generates reactive oxygen species, which oxidizes the methionine at position 35 of Aβ, and sequentially enhances its aggregation in intracellular and extracellular environments.

Impairments in sleep and circadian rhythms, such as difficulty in inducing and maintaining sleep, increased time spent napping, shifts in sleep/wakefulness patterns, reductions in rapid eye movement (REM) sleep, and slow-wave deficits during non-REM (NREM) sleep, have been increasingly reported in AD patients^20–22^. These disturbances are also noted to have a reciprocal association with cognitive impairments. Sleep alterations usually precede the early stage of AD. More than 65% of patients with AD or mild cognitive impairment have been reported to have at least one clinical sleep disorder^23^. Sleep impairments have also been reported to occur in model mice of familial AD ^24–29^. However, the sleep symptoms that occur in both AD patients and AD model mice vary substantially among individuals. Furthermore, it is still unclear which types of Aβ accumulate in specific brain regions and how they contribute to different sleep disorders. To address this, *in vivo* exogenous seeding models, in which specific types of Aβ are used as “seeds” and injected into animals to induce specific types of Aβ aggregates and fibrils, are a useful experimental technique. Seeding experiments enable the spatial and temporal onset of amyloidosis to be controlled. Numerous studies have performed seeding experiments in wild-type mice, AD model mice, rats, and non-human primates as hosts. Additionally, various types of synthesized Aβ, and extracts from the brains of AD model mice and AD patients have been used as seeds for administration; and it has been demonstrated that the effect of seeding is variable depending on the combination of the host and the administered agent^30,31^. Thus, most *in vivo* seeding studies have focused on the seeding potency and spreading ability -the capacity of aggregates to propagate and induce pathology in connected brain regions-of various Aβ species. To our knowledge, studies on whether exogenous Aβ administration causes pathological changes *in vivo*, particularly impairments of sleep/wakefulness states, are limited to date.

In this study, to investigate whether synthesized various Aβ species can directly affect sleep/wakefulness states and cortical oscillations, we bilaterally injected Aβ40 amyloid fibrils and Aβ42 amyloid fibrils into the brains of approximately 3-month-old wild-type mice, meaning they were not aged mice. In humans, approximately 90% of Aβ produced in human plasma and CSP is Aβ40, whereas the remaining Aβ species is primarily Aβ42. Although Aβ40 and Aβ42 differ by only two residues (Ile41 and Ala42) at the C-terminus, they have different molecular and pathological effects, including aggregation propensity, physicochemical properties, and cytotoxicity^6,32–36^. Although small Aβ oligomers have been shown to have strong neurotoxicity, amyloid fibrils in AD have been extensively studied owing to their localization in plaques, cytotoxicity, cross-seeding ability, and roles in disease progression^37–39^. Interestingly, previous studies have shown that Aβ42 amyloid fibrils are more toxic to neurons and trigger stronger neuroinflammatory responses than Aβ40 amyloid fibrils^40^. These differences in their deleterious effects may be linked to the distinct three-dimensional structures or polymorphisms of Aβ40 and Aβ42 amyloid fibrils^41–47^, similar to the polymorphism observed in α-synuclein amyloid fibrils observed in patients with Parkinson’s disease (PD)^48^.

In this study, we first aimed to clarify the differences between Aβ40 and Aβ42 in their aggregation behavior and amyloid fibril structures through *in vitro* experiments. We then aimed to elucidate whether the administration of a single type of Aβ amyloid fibril to wild-type mice could directly cause neurodegeneration or affect sleep/wakefulness states and cortical oscillations. To investigate these points, we injected Aβ40 and Aβ42 amyloid fibrils into the dentate gyrus (DG) of the hippocampus, which is one of the first regions to develop plaques in human AD patients^49^. We further assessed whether the resulting effects differ between Aβ40 and Aβ42 fibrils. Our results showed that both Aβ40 and Aβ42 formedβ-structured amyloid fibrils with distinct aggregation kinetics. Aβ42 aggregated into amyloid fibrils faster than Aβ40. Biophysical characterization experiments demonstrated that Aβ40 and Aβ42 developed into distinct amyloid fibrils with different physicochemical properties regarding their structure, secondary structure, and morphology. Our *in vivo* analyses demonstrated that the administration of Aβ42 into the hippocampus of mice substantially affected both their sleep/wakefulness state and cortical oscillations, whereas the administration of Aβ40 had minimal effect on their sleep/wakefulness state, but affected cortical oscillations. Neuronal death was only observed in the Aβ42-treated mice. These results suggest that Aβ40 and Aβ42 can induce distinct pathological changes *in vivo*, paving the way for the development of targeted treatments for each in AD.

## Results

### Characterization of distinct amyloidogenesis characteristics of Aβ40 and Aβ42

The detection of amyloid formation and the differentiation of various amyloid fibril types can be performed using various biophysical approaches, including fluorescence, nuclear magnetic resonance (NMR), and microscopy^3,36,50,51^. We first performed the thioflavin T (ThT) fluorescence assay to investigate the aggregation behaviors of Aβ40 and Aβ42 (Fig. 1a–d), as ThT is a fluorescent dye commonly used to monitor amyloid formation^52^. Regarding Aβ40 amyloid formation, ThT fluorescence intensity increased after a lag phase of approximately 2 h for nucleation, with an elongation rate constant of approximately 12 h^-1^, and reached a plateau at about 3 h (Fig. 1a). This sigmoidal kinetic profile indicates nucleation-dependent amyloid formation, which is a process that is usually observed in protein amyloidogenesis^4,6,51,53,54^. Aβ42 showed even faster nucleation-dependent amyloid formation, with a shorter lag phase of approximately 0.1 h, and an elongation rate constant of about 31 h^-1^, which was higher than that for Aβ40 (Fig. 1a–c). Of note is that the maximum ThT intensity of Aβ42 was markedly lower than that of Aβ40, which can be explained by the formation of polymorphic amyloid structures by Aβ42 (Fig. 1d), as ThT binding is specific to fibrillar structures^55^. The aggregation of Aβ40 and Aβ42 was further supported by the results of ^1^H NMR spectroscopy (Fig. 1e and f), based on NMR peak intensity changes. We have previously shown that NMR signals significantly decrease following amyloid formation owing to the slow tumbling of molecules with a large molecular weight^4,6,56^. Before incubation, the ^1^H NMR spectra of both Aβ40 and Aβ42 showed a narrow distribution with peaks between approximately 7.2 and 8.5 parts per million (ppm), indicating the presence of monomers with random-coil structures^4,51^. However, the substantial disappearance of these peaks after incubation confirmed that almost all Aβ monomers had formed amyloid fibrils with a large molecular weight.

**Figure 1.**
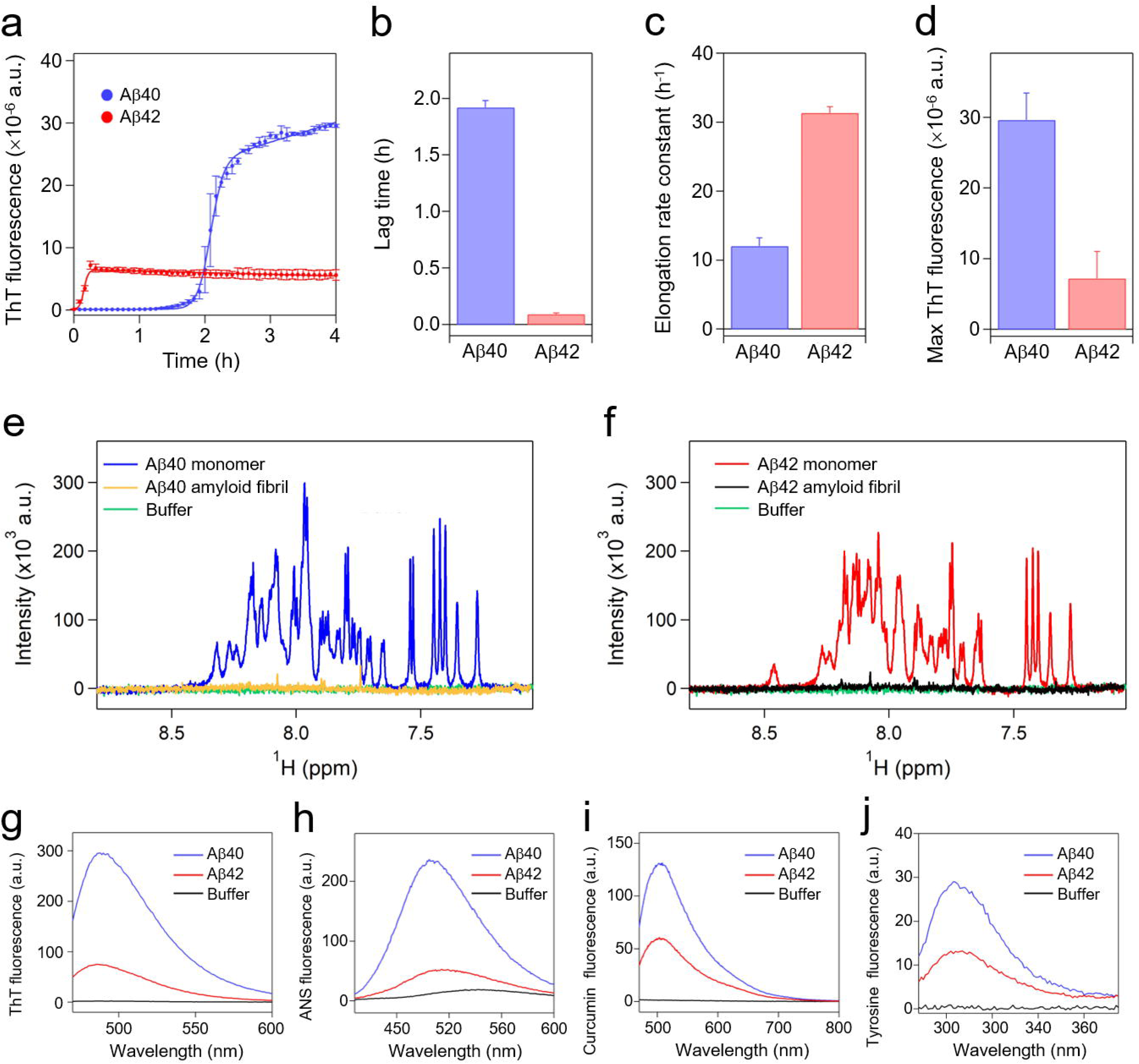
Molecular characterization of. **A**β**40 and A**β**42 amyloid fibrils** (a) Amyloid formation of Aβ40 and Aβ42 monitored using ThT fluorescence. Solid lines represent the fit curves. Lag times (b), elongation rate constants (c), and maximum ThT fluorescence intensities (d) of Aβ40 and Aβ42 amyloidogenesis. Average values calculated from three separate samples are shown, with error bars indicating the standard deviation. One-dimensional proton NMR spectra of both monomers and amyloid fibrils of Aβ40 (e) and Aβ42 (f). Fluorescence spectra of ThT (g), ANS (h), and curcumin (i), in the presence of Aβ40 and Aβ42 amyloid fibrils. (j) Tyrosine fluorescence spectra of Aβ40 and Aβ42 amyloid fibrils.

To further investigate the polymorphic nature of Aβ40 and Aβ42 amyloid fibrils, we analyzed the post-incubation samples using fluorescence dye-binding assays, intrinsic tyrosine fluorescence, circular dichroism (CD) spectroscopy, atomic force microscopy (AFM), and transmission electron microscopy (TEM) (Fig. 1g–j and Fig. 2a–h). As shown in Fig. 1g–i, dye-binding assays demonstrated distinct fluorescence spectra for Aβ40 and Aβ42 species after incubation. The ThT fluorescence spectrum of Aβ40 amyloid fibrils showed a red shift in the wavelength of maximum fluorescence, and increased intensity compared with Aβ42 amyloid fibrils. (Fig. 1g). 8-anilino-1-naphthalenesulfonic acid (ANS) is a fluorescence dye commonly used to detect exposed hydrophobic areas in proteins^57^. ANS fluorescence analysis demonstrated that Aβ40 amyloid fibrils demonstrate a blue shift in maximum wavelength, with increased intensity compared with Aβ42 amyloid fibrils, indicating that the hydrophobic regions of Aβ40 amyloid fibrils are more exposed than those of Aβ42 amyloid fibrils (Fig. 1h). Curcumin is widely used to study amyloid fibrils because it binds to β-sheet-rich structures, leading to enhanced fluorescence that facilitates the visualization and characterization of amyloid fibrils^51,58^. Curcumin fluorescence also showed a difference in maximum intensity between Aβ40 and Aβ42 amyloid aggregates (Fig. 1i). In addition, Aβ40 and Aβ42 amyloids demonstrated different fluorescence intensities of the intrinsic tyrosine (Tyr10), suggesting that the tyrosine resides of Aβ40 and Aβ42 amyloid fibrils are located in different environments (Fig. 1j). All these tinctorial results confirm that Aβ40 and Aβ42 amyloid fibrils have different characteristics, even when formed under the same condition.

**Figure 2.**
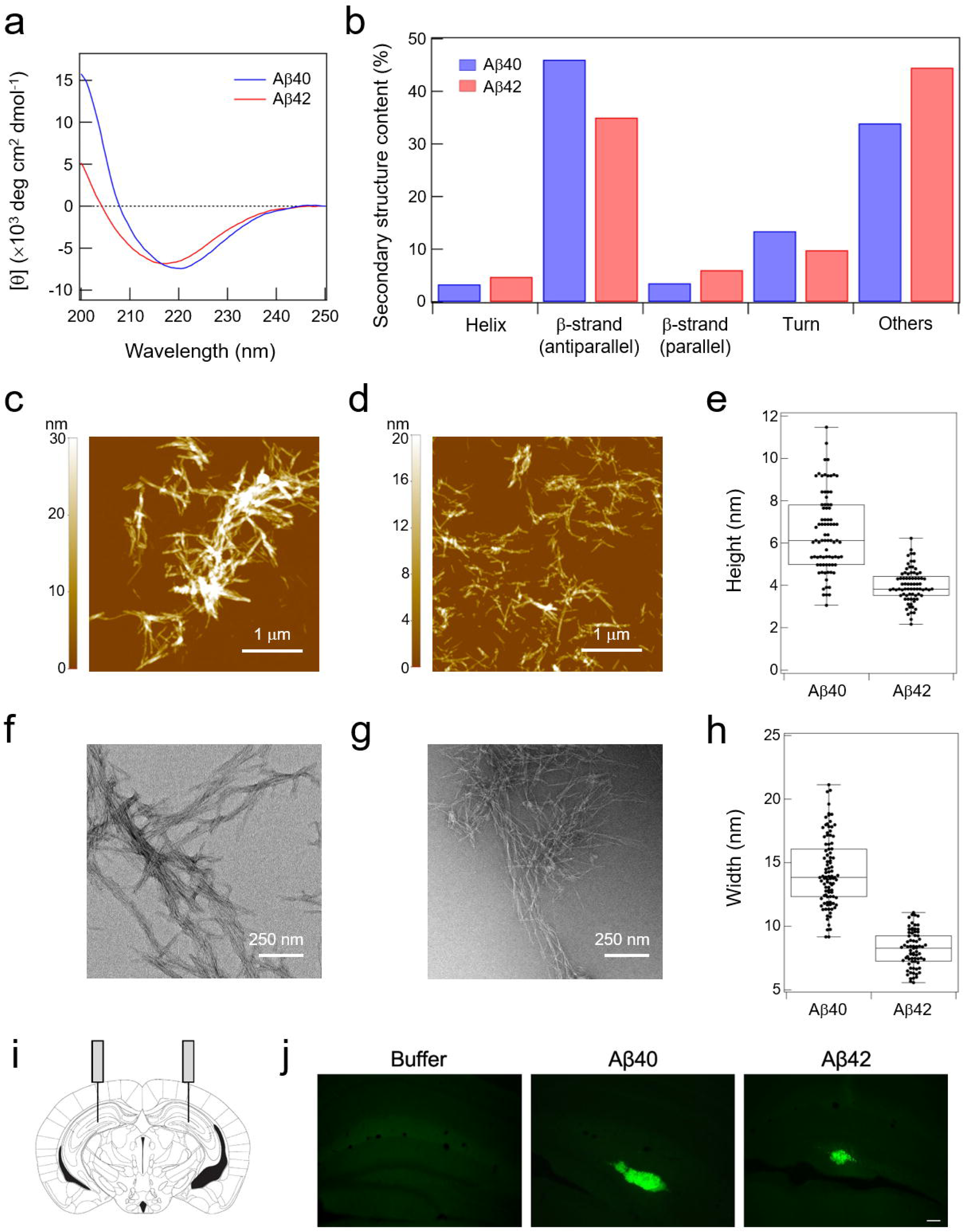
Differences in the characteristics of Aβ40 and Aβ42 amyloid fibrils, and their deposition in the hippocampal region. (a) Far-UV CD spectra of Aβ40 and Aβ42 amyloid fibrils. (b) Secondary structure contents of Aβ40 and Aβ42 amyloid fibrils estimated using far-UV CD spectra and the BeStSel algorithm^61^. Others include all non-helical, non β-sheet, and non-turn structures, primarily characterized as disordered structures. AFM (c and d) and TEM (f and g) images of Aβ40 and Aβ42 amyloid fibrils. The white scale bars in the AFM and TEM images represent 1 μm and 250 nm, respectively. Height (e) and width (h) distributions of Aβ40 and Aβ42 amyloid fibrils were calculated using AFM and TEM images, respectively. Each sample set contains approximately 80 points. Box plots indicate the interquartile range, and error bars represent one standard deviation. (i) Schematic drawing showing the position of Aβ injection into the mouse hippocampus. (j) Immunohistochemical analysis demonstrated that anti-fibril Aβ immunostaining (green) is observed in the hippocampus of mice administered with Aβ40 and Aβ42. Brain sections of buffer-treated mice (left), Aβ40-treated mice (middle), and Aβ42-treated mice (right). Scale bar: 100 μm

### Differences in A**β**40 and A**β**42 amyloid fibril deposition in the brain

Consistent with the results of fluorescence analyses, CD spectroscopy demonstrated distinct spectra of Aβ40 and Aβ42 amyloid fibrils, showing negative peaks at approximately 220 nm and 217 nm, respectively (Fig. 2a). As far-ultraviolet (UV) CD spectra reflect the secondary structure of proteins and peptides^14,53,59,60^, the distinct far-UV CD spectra of Aβ40 and Aβ42 amyloid fibrils suggested that they have different secondary structures. Secondary structure analysis based on the CD results, using the BeStSel algorithm^61^, further demonstrated that Aβ40 and Aβ42 amyloid fibrils differ in their secondary structural composition (Fig. 2b). Aβ40 amyloid fibrils contain a higher proportion of antiparallel β-structures and a lower proportion of disordered regions compared with Aβ42 amyloid fibrils. Various atomic-level structures of Aβ40 and Aβ42 amyloid fibrils have also been reported^43,62–64^. To obtain more information on the structural differences of these fibrils, we performed microscopic analyses. AFM images demonstrated that Aβ40 amyloid fibrils are longer and tend to cluster, whereas Aβ42 amyloid fibrils are shorter and more dispersed (Fig. 2c and d). Further imaging analyses indicated that the height of Aβ40 amyloid fibrils (∼6.5 nm) was greater than that of Aβ42 amyloid fibrils (∼4.0 nm) (Fig. 2e). TEM micrographs demonstrated that Aβ40 amyloid fibrils are thicker than Aβ42 amyloid fibrils (Fig. 2f and g). The average width was approximately 14.3 nm and 8.2 nm for Aβ40 and Aβ42 amyloid fibrils, respectively (Fig. 2h). Taken all together, differences in the kinetics of amyloid formation, surface and secondary structures, and physicochemical and morphological properties demonstrate that Aβ40 and Aβ42 form amyloid fibrils with distinct characteristics that may have different biological and pathological effects on neurons and individuals.

Next, we investigated whether Aβ deposition occurs upon the exogenous administration of purified Aβ40 amyloid fibrils and Aβ42 amyloid fibrils to the hippocampal parenchyma. A previous study reported that no Aβ deposition was observed upon the administration of dilute Aβ-containing brain extracts (estimated Aβ concentration: 1 to 10 ng/μL) from human patients with Alzheimer’s disease or APP transgenic mice, a model of Alzheimer’s disease^65^. Injection of synthetic Aβ at concentrations 100 to 1,000 times that of Aβ in APP23 transgenic mouse brain extracts resulted in Aβ deposition. Therefore, we injected 1.5 μL of 100 μM Aβ40 and Aβ42 (600–700 ng/mL) into the brains of mice in this study. We bilaterally injected Aβ40 fibrils, Aβ42 fibrils, and buffer as a control into the DG regions of the hippocampus of approximately 3-month-old wild-type mice (Fig. 2i). Immunohistochemical analyses using the anti-amyloid fibril antibody revealed that the administration of Aβ40 and Aβ42 induces the formation of Aβ deposition in the DG (Fig. 2j). On the other hand, no Aβ deposition was detected in the case of buffer administration (Fig. 2j). In addition, glial cell activation was observed in the depositions formed by Aβ40 administration (Extended Fig. 1). Interestingly, reactive astrocytes accumulated surrounding the Aβ depositions, and reactive microglia accumulated overlapping with Aβ deposition.

### Injection of A**β**42 amyloid fibrils into the hippocampus decreases total time of REM sleep

To assess whether bilateral injection of Aβ40 amyloid fibrils and Aβ42 amyloid fibrils into the hippocampus of wild-type mice affects their sleep/wakefulness state, 24-hour electroencephalogram (EEG) and electromyogram (EMG) recordings were performed using freely moving mice. To observe progressive changes, continuous recordings were carried out from day 3 to day 14 after Aβ fibril administration, and averages were calculated every three days (Fig. 3a). As marked alterations were observed in the data from day 12 to day 14 after Aβ fibril administration (Day12–14), we focused on Day12–14 data thereafter, but data on the sleep/wakefulness state at earlier timepoints are shown in Extended Figures 2 to 4.

**Figure 3.**
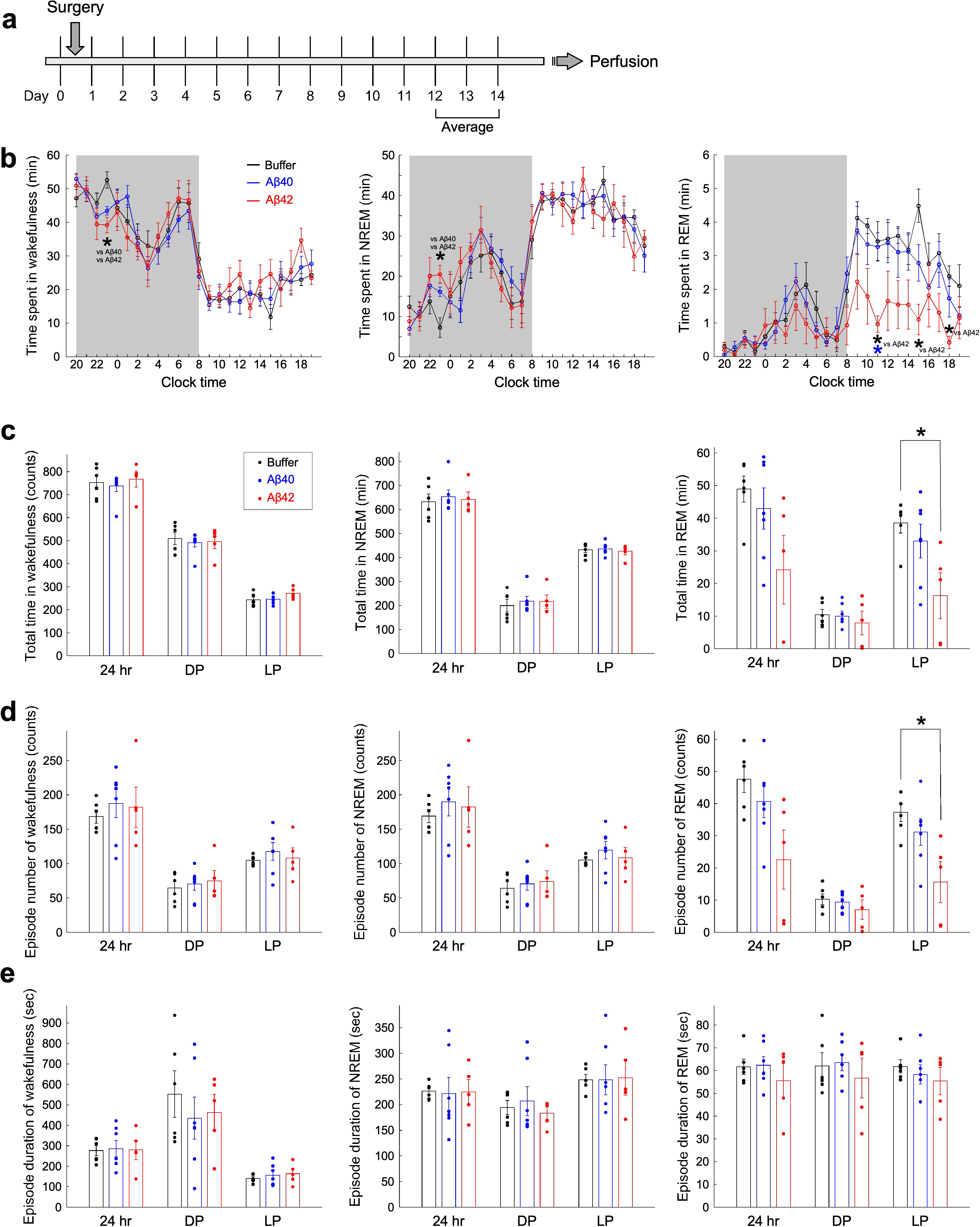
Sleep/wakefulness patterns of Aβ40-treated and Aβ42-treated mice, 12 to 14 days after Aβ administration. (a) Schematic drawing showing the experimental protocol. To compensate for daily variations, three-day averages were calculated. (b) The hourly amount of wakefulness (left), NREM sleep (middle), and REM sleep (right) during 24 hours. The average of days 12 to 14 after injection are shown. The gray background indicates the dark period. Wakefulness: at 23:00, Kruskal–Wallis: F(2,15) = 10.9, *p* < 0.01; post hoc Steel-Dwass test, buffer vs Aβ40, *p* = 0.0182; buffer vs Aβ42, *p* = 0.0286. NREM: at 23:00, Kruskal–Wallis: F(2,15) = 10.5, *p* < 0.01; post hoc Steel-Dwass test, buffer vs Aβ40, *p* = 0.0274; buffer vs Aβ42, *p* = 0.0286. REM: at 11:00, Kruskal–Wallis: F(2,15) = 10.3, *p* < 0.01; post hoc Steel-Dwass test, buffer vs Aβ42, *p* = 0.0217; Aβ40 vs Aβ42, *p* = 0.0155; at 15:00, Kruskal–Wallis: F(2,15) = 9.5, *p* < 0.01; post hoc Steel-Dwass test, buffer vs Aβ42, *p* = 0.0170; at 18:00, Kruskal–Wallis: F(2,15) = 7.8, *p* < 0.05; post hoc Steel-Dwass test, buffer vs Aβ42, *p* = 0.0167. (c) Total amount of wakefulness (left), NREM sleep (middle), and REM sleep (right). REM: LP, Kruskal–Wallis: F(2,15) = 6.7, *p* < 0.05; post hoc Steel-Dwass test, buffer vs Aβ42, *p* = 0.0286. (d) Number of episodes of wakefulness (left), NREM sleep (middle), and REM sleep (right). REM: LP, Kruskal–Wallis: F(2,15) = 8.4, *p* < 0.05; post hoc Steel-Dwass test, buffer vs Aβ42, *p* = 0.0281. (e) Mean episode duration of wakefulness (left), NREM sleep (middle), and REM sleep (right). DP, dark period; LP, light period. Values are presented as means ± standard error of the mean (SEM); *, *p* < 0.05.

**Figure 4.**
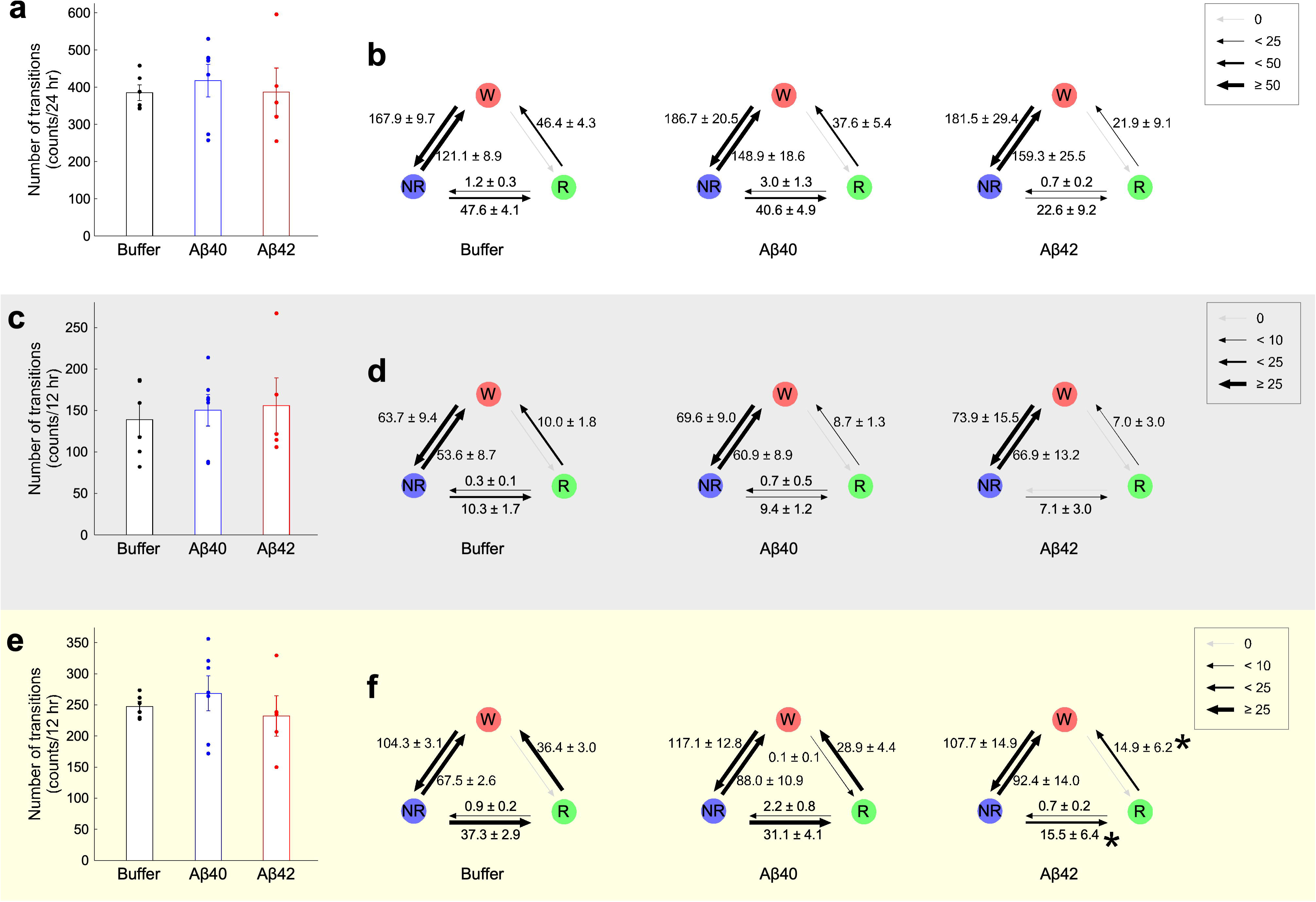
Frequency of sleep/wakefulness stage transitions in Aβ40-treated and Aβ42-treated mice. (a, c, and e) Number of transitions between all states during 24 hours (a), the DP (c), and the LP (e) on days12 to 14 after Aβ administration. (b, d, and f) Number of transitions between each sleep/wakefulness state during 24 hours (b), the DP (d), and the LP (f). Left, buffer-treated mice; middle, Aβ40-treated mice; left, Aβ42-treated mice. From NREM to REM: Kruskal–Wallis: F(2,15) = 8.4, *p* < 0.05; post hoc Steel-Dwass test, buffer vs Aβ42, *p* = 0.0281. From REM to wakefulness: Kruskal–Wallis: F(2,15) = 7.8, *p* < 0.05; post hoc Steel-Dwass test, buffer vs Aβ42, *p* = 0.0286. Gray background indicates dark period. Yellow background indicates light period. NR, NREM sleep; R, REM sleep; W, wakefulness. Values are presented as means ± SEM; *, *p* < 0.05

Hourly amounts of sleep/wakefulness during 24 hours showed that wakefulness was significantly reduced and NREM sleep was significantly increased in Aβ40-treated mice and Aβ42-treated mice at one point in the dark period (DP), which corresponds to the active phase for mice, compared with buffer-treated mice (Fig. 3b). There were no significant differences in total time, episode number, or episode duration of wakefulness and NREM sleep during the 24 hours, the DP, and the light period (LP) (Fig. 3c–e) among the mice. In contrast, a remarkable change in REM sleep was observed, particularly during the LP. The hourly amount of REM sleep during the LP was significantly reduced in Aβ42-treated mice compared with buffer-treated mice (Fig. 3b). The total time of REM sleep during the LP in Aβ42-treated mice significantly decreased compared with buffer-treated mice. There were no significant changes in episode duration of REM sleep, whereas the number of episodes of REM sleep was significantly decreased in Aβ42-treated mice (Fig. 3d, e). The transition frequencies of all sleep/wakefulness states were not different among the mice during the 24 hours, the DP, and the LP (Fig. 4a, c, e). Next, the frequencies of transition to each state were further analyzed in detail (Fig. 4b, d, f). During the LP, the number of transitions from NREM sleep to REM sleep and from REM sleep to wakefulness was significantly decreased in Aβ42-treated mice compared with buffer-treated mice, but not in Aβ40-treated mice (Fig. 4f). These results suggest that Aβ42-treated mice impair REM sleep, with a decrease in episode number resulting in a decrease in total REM sleep time.

### Injection of A**β**40 and A**β**42 amyloids into the hippocampus alters cortical oscillations

Next, alterations in the cortical oscillations of mice in each sleep/wakefulness state were analyzed based on their EEG. The EEG power of each frequency bin was represented as a normalized power, by dividing by the total EEG power over all frequency bins (0–20 Hz) (Fig. 5). Fig. 5 represents the data of Day12–14. Neither the Aβ40-treated or Aβ42-treated mice showed any changes in normalized EEG during NREM sleep (Fig. 5b, e, h, k). On the other hand, substantial changes were observed in wakefulness and REM sleep, as shown in the power spectra (Fig. 5a, c). During wakefulness, normalized delta power significantly increased in Aβ40-treated mice compared with buffer-treated mice, but not in Aβ42-treated mice (Fig 5d). Normalized theta power significantly decreased in Aβ40-treated mice and Aβ42-treated mice during 24 hours and the LP (Fig. 5g). Theta/delta ratio was reduced only in Aβ40-treated mice (Fig. 5j). During REM sleep, Aβ42-treated mice had significantly increased normalized delta power and decreased normalized theta power compared with buffer-treated mice (Fig. 5f, i). Aβ40-treated mice had significantly decreased normalized theta power during only the LP compared with buffer-treated mice (Fig. 5i). The theta/delta ratio during REM sleep was significantly reduced only in Aβ42-treated mice (Fig. 5l). These results interestingly suggest that Aβ40 and Aβ42 amyloid depositions have different effects on cortical oscillations of mice. An increase in theta power during REM sleep is a characteristic of REM sleep. We therefore next investigated whether REM sleep induced by Aβ administration exhibits rapid eye movement, which is the origin of the name “REM” sleep, even if theta power is decreased. As shown in Extended Movie 1, despite a reduction in the theta component, rapid eye movement was observed. Data of cortical oscillations from 3 to 11 days after Aβ administration are shown in Extended Figures 5 to 7. Analysis of changes in sleep/wakefulness states found that significant differences in REM sleep were also observed on Days 6 to 8 (Extended Figs. 2–4), but these differences were more pronounced on Day 12–14. However, a significant difference in EEG was observed from day 6 after Aβ administration (Extended Figs. 5–7). This suggests that cortical oscillation changes precede sleep/wakefulness changes.

**Figure 5.**
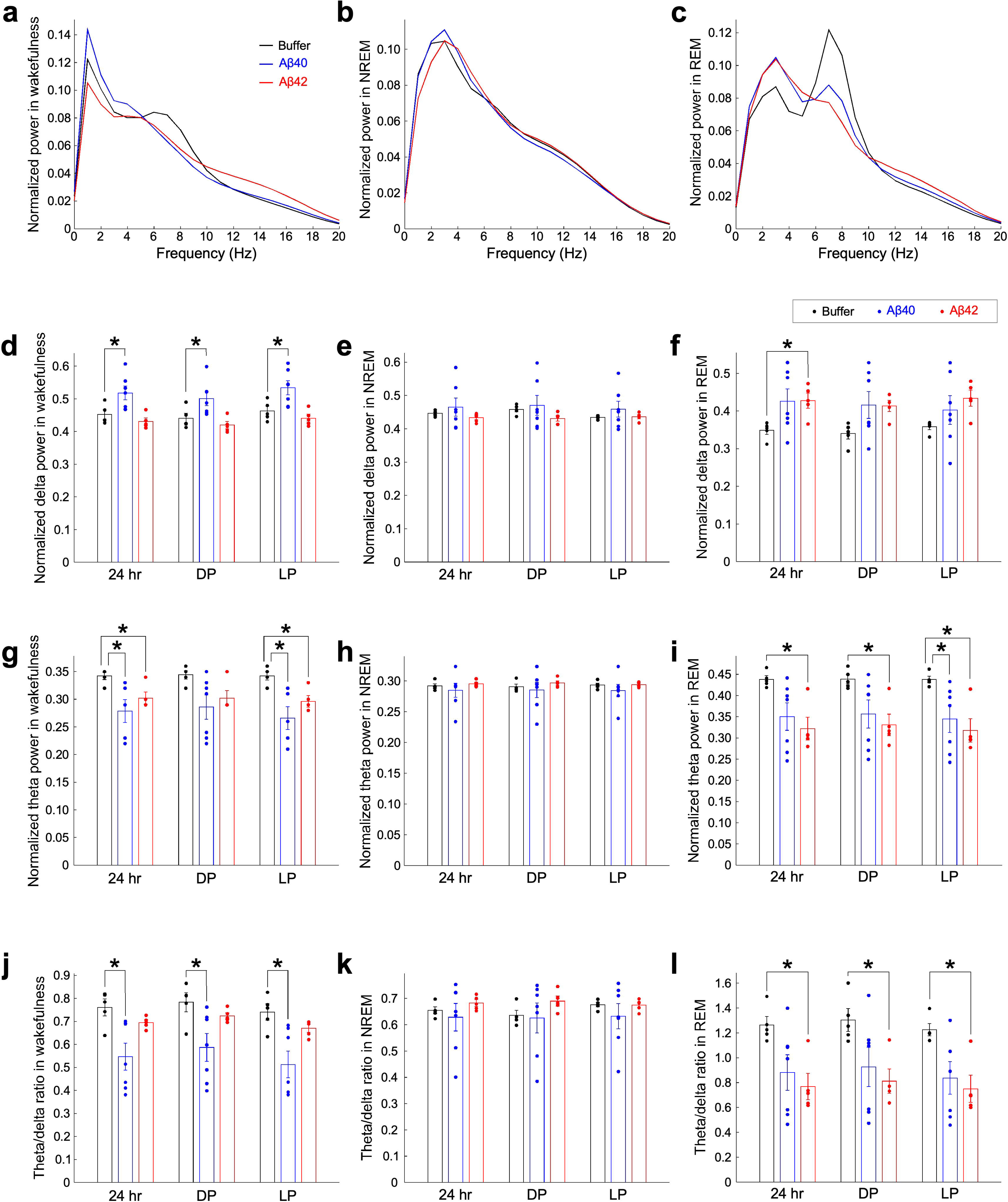
EEG power spectra during wakefulness, NREM, and REM sleep in Aβ40-treated and Aβ42-treated mice. (a–c) Average EEG power spectra during wakefulness (a), NREM sleep (b), and REM sleep (c), 12 to 14 days after Aβ administration. (d–f) Comparison of normalized delta (1–5 Hz) power during wakefulness (d), NREM sleep (e), and REM sleep (f). Wakefulness: 24 hours, Kruskal–Wallis: F(2,14) = 11.0, *p* < 0.01; post hoc Steel-Dwass test, buffer vs Aβ40, *p* = 0.0188; DP, Kruskal–Wallis: F(2,14) = 9.8, *p* < 0.01; post hoc Steel-Dwass test, buffer vs Aβ40, *p* = 0.0291; LP, Kruskal–Wallis: F(2,14) = 8.5, *p* < 0.05; post hoc Steel-Dwass test, buffer vs Aβ40, *p* = 0.0244. REM: 24 hours, Kruskal–Wallis: F(2,14) = 6.4, *p* < 0.05; post hoc Steel-Dwass test, buffer vs Aβ42, *p* = 0.0313. (g–i) Comparison of normalized theta (6–10 Hz) power during wakefulness (g), NREM sleep (h), and REM sleep (i). Wakefulness: 24 hours, Kruskal–Wallis: F(2,14) = 7.8, *p* < 0.05; post hoc Steel-Dwass test, buffer vs Aβ40, *p* = 0.0364, buffer vs Aβ42, *p* = 0.0480; LP, Kruskal–Wallis: F(2,14) = 8.5, *p* < 0.05; post hoc Steel-Dwass test, buffer vs Aβ40, *p* = 0.0186, buffer vs Aβ42, *p* = 0.0414. REM: 24 hours, Kruskal–Wallis: F(2,14) = 7.6, *p* < 0.05; post hoc Steel-Dwass test, buffer vs Aβ42, *p* = 0.0307; DP, Kruskal–Wallis: F(2,14) = 7.6, *p* < 0.05; post hoc Steel-Dwass test, buffer vs Aβ42, *p* = 0.0398; LP, Kruskal–Wallis: F(2,14) = 8.7, *p* < 0.05; post hoc Steel-Dwass test, buffer vs Aβ40, *p* = 0.0247, buffer vs Aβ42, *p* = 0.0287. (j–l) Ratio of theta to delta power during wakefulness (j), NREM sleep (k), and REM sleep (l). Wakefulness: 24 hours, Kruskal–Wallis: F(2,14) = 7.4, *p* < 0.05; post hoc Steel-Dwass test, buffer vs Aβ40, *p* = 0.0476; DP, Kruskal–Wallis: F(2,14) = 6.8, *p* < 0.05; post hoc Steel-Dwass test, buffer vs Aβ40, *p* = 0.0481; LP, Kruskal–Wallis: F(2,14) = 9.0, *p* < 0.05; post hoc Steel-Dwass test, buffer vs Aβ40, *p* = 0.0385. REM: 24 hours, Kruskal–Wallis: F(2,14) = 6.9, *p* < 0.05; post hoc Steel-Dwass test, buffer vs Aβ42, *p* = 0.0313; DP, Kruskal–Wallis: F(2,14) = 8.7, *p* < 0.05; post hoc Steel-Dwass test, buffer vs Aβ42, *p* = 0.0422; LP, Kruskal–Wallis: F(2,14) = 7.5, *p* < 0.05; post hoc Steel-Dwass test, buffer vs Aβ42, *p* = 0.0218. DP, dark period; LP, light period. Values are presented as means ± SEM; *, *p* < 0.05.

### Injection of A**β**42 amyloid fibrils induces neuronal loss in the hippocampus

Finally, we considered that the impairment of REM sleep, and cortical oscillations during wakefulness and REM sleep upon the injection of Aβ into the hippocampus may be owing to the loss of hippocampal neurons. To address this, NeuroTrace Nissl staining was performed (Fig. 6). Images of NeuroTrace Nissl staining showed that the administration of Aβ, particularly Aβ42 into the mouse hippocampus resulted in less staining compared with buffer-treated mice, implying neuronal loss (Fig. 6a). To quantify this, the entire hippocampus was used as the region of interest, and the percentage area over threshold was calculated based on fluorescence intensity (Fig. 6b). We found a significant decrease in the percentage area over threshold in Aβ42-treated mice compared with buffer-treated and Aβ40-treated mice.

**Figure 6.**
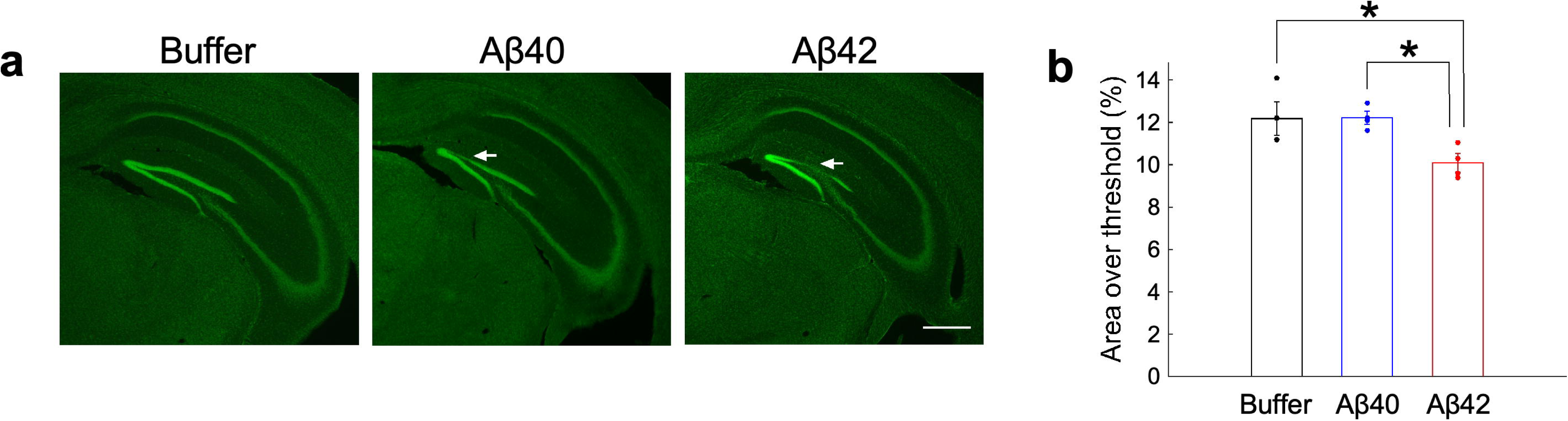
Hippocampal neuronal loss in Aβ42-treated mice. (a) Nissl staining (green) demonstrating hippocampal neuronal loss (arrows) in Aβ-treated mice. Representative brain sections of buffer-treated mice (left), Aβ40-treated mice (middle), and Aβ42-treated mice (right). Scale bar: 500 μm. (b) Percentage area over threshold calculated from the Nissl-stained hippocampal sections of the various mice. One-way analysis of variance (ANOVA): F(2, 9) = 6.5, *p* < 0.05; post hoc Fisher’s protected least significant difference test, buffer vs Aβ42, *p* = 0.0130, Aβ40 vs Aβ42, *p* = 0.0117. Values are presented as means ± SEM; *, *p* < 0.05.

## Discussion

In the present study, we analyzed the alterations in sleep/wakefulness states and cortical oscillations caused by the administration of single types of Aβ amyloid fibrils into the DG of the hippocampus of wild-type mice. Our study demonstrated that the administration of Aβ42 amyloid fibrils significantly reduced time spent in REM sleep during the LP, together with the suppression of REM sleep episode occurrence, whereas the administration of Aβ40 amyloid fibrils had minimal effects on the sleep/wakefulness state. On the other hand, cortical oscillations were observed upon both Aβ40 and Aβ42 administration, but in a different manner. Aβ40-treated mice showed a significant increase in the delta component and decrease in the theta component during wakefulness compared with buffer-treated mice. In contrast, Aβ42-treated mice showed significantly reduced delta components during both wakefulness and REM sleep compared with buffer-treated mice. These results indicate that a single administration of Aβ can alter the sleep/wakefulness state and cortical oscillations *in vivo*, and the effects vary depending upon the type of Aβ administered.

The distinct pathological effects of Aβ40 and Aβ42 amyloid fibrils on mouse brains can be attributed to differences in their surface properties, structures, and morphology. Previous studies demonstrated strong associations between the structural properties of biomolecular assemblies and their biological functions and toxicity^66–69^. In addition, distinct structural states of α-synuclein amyloid fibrils in PD formed under various conditions demonstrated varying cytotoxicity^48^. Our far-UV CD analyses showed that Aβ42 amyloid fibrils have a higher proportion of disordered regions than Aβ40 amyloid fibrils. It has been suggested that disordered regions of amyloid fibrils play a key role in enabling amyloid fibrils to disrupt cell membranes and RNA, leading to impaired cellular quality control systems^70^. Thus, our structure-based results are consistent with these previous findings. In addition to conformational effects, the size of amyloid fibrils also plays a crucial role in their neurotoxic effects. Smaller amyloid fibrils have been shown to have a stronger capacity to disrupt cell membranes and act as seeds for further amyloidogenesis^71,72^. Consistent with this, AFM and TEM imaging demonstrated that Aβ42 fibrils are smaller in both width and height than Aβ40 fibrils, suggesting that the shorter size of Aβ42 fibrils may increase their toxic properties. Taken together, these structural and morphological differences may explain why Aβ42 amyloid fibrils have a stronger negative effect on sleep and wake states, as well as on cortical oscillations, compared with Aβ40 amyloid fibrils.

### Neuropathological changes following a single administration of A**β** fibrils

To identify which type of Aβ accumulation in which brain region leads to the various symptoms of AD, it is vital to administer synthetic Aβ to specific brain regions and assess its effect on neuropathology. A previous study by Meyer-Luehmann et al. demonstrated that seeding with Aβ40 amyloid fibrils, Aβ42 amyloid fibrils, and Aβ42 oligomers was insufficient to induce amyloidosis in APP transgenic mice^65^. Additionally, another study demonstrated that a single administration of either Aβ40 monomers or Aβ43 monomers had little effect on spatial learning or synaptic transmission and plasticity^73^. Recent studies, however, have reported that the administration of a high amount of synthetic Aβ fibrils induces amyloidosis in mice^74,75^. In the present study, we were able to observe the formation of plaques, which are localized accumulation of Aβ amyloid fibrils, and the subsequent development of amyloidosis, a broader pathological condition caused by amyloid deposition, in the brains of wild-type mice. This was achieved by injecting Aβ fibrils at amounts greater than 100 times those used in the study Meyer-Luehmann et al..

Previous studies using APP-overexpressing transgenic mice have shown that the accumulation of insoluble Aβ has minimal effects on neuronal loss^76,77^, indicating that a species other than insoluble fibrillar Aβ may underlie neuronal loss. In the present study, however, we observed neuronal loss in Aβ42-treated mice, implying that insoluble fibrillar Aβ can directly contribute to neurotoxicity. In contrast, Aβ40 fibrils had minimal effect on neurotoxicity and neuronal loss. Similar neurotoxic effects have also been reported for the fibrillar forms of other amyloidogenic proteins, such as α-synuclein and tau which are involved in the pathogenesis of PD^78,79^. Of note, the progression of Aβ plaques was not observed in our study, suggesting that a single administration of Aβ amyloid fibrils is insufficient to induce plaque progression. This indicates the potential involvement of additional cofactors or specific microenvironmental factors in the induction of plaque progression. Previous studies have highlighted that alterations in neuronal lipid metabolism, microglial activation, and innate immune responses play a crucial role in this process^80–82^. These findings suggest that Aβ plaque progression is a complex process involving multiple molecular pathways. Future research is required to identify the specific molecular and environmental conditions that facilitate plaque progression as this could provide valuable insights into the pathogenesis of AD.

### REM sleep impairments in A**β**42-treated mice

In this study, we showed that Aβ42 administration into the hippocampus impairs REM sleep, particularly during the LP, although Aβ40 administration has minimal effect on the sleep/wakefulness state. The total time of REM sleep significantly decreased, together with a reduction in REM sleep episode occurrence. Previous studies reported that various AD model mice show REM sleep impairments, i.e., reduced total time and reduced episode duration of REM sleep^24–26,28,83,84^. Based on these studies, it can be concluded that the accumulation of Aβ42 in the hippocampus may have a specific effect on REM sleep. AD model mice often show impairment not only in REM sleep but also in NREM sleep. There is a possibility that effects of Aβ42 on the sleep/wakefulness state, excluding REM sleep, may be owing to effects on brain areas other than the hippocampus. According to previous reports, disruption of the sleep/wakefulness state in AD patients results from damage to cholinergic neurons in the basal forebrain^85–87^. However, our present study suggests that this may not be the case. Aβ42 administration to the hippocampus induced hippocampal neuronal loss, but there has been no evidence to date that hippocampal neurons are responsible for controlling sleep/wakefulness states. In addition, the possible contribution of glial cells cannot be ignored. As we showed in the present study, reactive astrocytes and activated microglia enhance Aβ deposition. Recently, we demonstrated that the sleep/wakefulness state can be modulated by the chemogenetic activation of hippocampal astrocytes in mice^88^. On the other hand, microglia have also been implicated in sleep/wakefulness regulation, and it is well known that they are activated during sleep deprivation^89–91^. Further studies are required to clarify the hippocampal mechanism underlying REM sleep impairments.

### Changes in cortical oscillations in A**β**40-treated and A**β**42-treated mice

Although the administration of Aβ40 into the hippocampus had minimal effects on the sleep/wakefulness state, cortical oscillations were affected; i.e., the delta component increased, and the theta component decreased during wakefulness. This effect was similar to that observed in Aβ42-treated mice. In contrast, Aβ42 administration significantly affected cortical oscillations during both REM sleep and wakefulness. Neither Aβ40-treated mice nor Aβ42-treated mice were affected during NREM sleep. How Aβ40 administration alters cortical oscillations without causing hippocampal neuronal loss remains unclear. One possibility is synaptic alterations. Aβ has been reported to impair N-methyl-D-aspartate receptor-dependent long-term potentiation in the hippocampus^92–96^. This might be associated with oscillatory activity changes in the brain. Both Aβ40 and Aβ42 administration had different effects on the cortical oscillations of mice, indicating that their toxicity could be mediated by different targets^97,98^. On the other hand, the administration of Aβ40 and Aβ42 had minimal effect on cortical oscillations during NREM sleep. It has been reported that cortical oscillations during NREM sleep are substantially affected in most AD model mice^25,83,99^. Compared with older individuals who do not have AD, AD patients were reported to have a decreased level of slow-wave activity during slow-wave sleep^100^. These symptoms are different from our present results, indicating that cortical oscillations during NREM sleep are important for Aβ accumulation in brain regions other than the hippocampus, such as the cortex and the thalamus, which are brain region crucial for the generation of cortical oscillations during NREM sleep^101^. Moreover, Aβ42-treated mice showed a decrease in theta components during REM sleep, which play a crucial role in memory consolidation^102^, indicating that the reduction in theta oscillations may also contribute to the progression of dementia.

The advantage of this study is that the experiments were performed on 3-month old adult wild-type mice, meaning they were not aged mice. AD model mice typically start to show *in vivo* symptoms after 6 months of age. It has been reported that old age affects sleep, and cortical oscillation impairments occur in wild-type mice at 6 months of age^103^. Therefore, we cannot conclude from AD model mice whether alterations in the sleep/wakefulness state and cortical oscillations are caused by old age or by AD. Our present study overcomes this problem. Another advantage of our study is that the effects of purely exogenous Aβ could be analyzed, as the Aβ was administered to wild-type mice. The type of Aβ administered and the brain region could be controlled, and thus their specific effects could be observed. As mentioned above, we demonstrated that Aβ42 had more severe pathological effects *in vivo* compared with Aβ40, meaning more toxicity. In a further study, we plan to repeat similar Aβ seeding experiments to investigate in detail the differences in symptoms when various types of Aβ are administered to different brain regions. As a result, prodromal symptoms of AD can be categorized, and clues towards the early detection of AD will be identified.

**Extended Figure 1. Aβ deposition induces the activation of astrocytes and microglia**

(a) Immunohistochemical analysis demonstrating reactive astrocyte accumulation around Aβ40 depositions in the hippocampus. Scale bar: 200 μm. The bottom row shows higher magnification images of the square regions in the top row. Green, anti-fibril Aβ immunoreactivity (IR); red, anti-glial fibrillary acidic protein (GFAP) immunoreactive cells; blue, 4’,6-diamidino-2-phenylindole (DAPI). Scale bar: 50 μm. (b) Immunohistochemical analysis demonstrating the colocalization of activated microglia with Aβ40 plaques in the hippocampus. Scale bar: 200 μm. The bottom row shows higher magnification images of the square regions in the top row. Green, anti-fibril Aβ immunoreactivity; red, anti-ionized calcium-binding adapter molecule 1 (Iba1) immunoreactive cells; blue, DAPI. Scale bar: 50 μm

**Extended Figure 2. Sleep/wakefulness patterns from 3 days to 11 days after Aβ injection**

(a) Schematic drawing showing the experimental protocol. To compensate for daily variations, three-day averages were calculated. (b–d) The hourly amount of wakefulness (b), NREM sleep (c), and REM sleep (d) over 24 hours. The graphs show the average of days 3 to 5 (left), 6 to 8 (middle), and 9 to 11 (right) after injection. Wakefulness: Day6–8, at 4:00, Kruskal–Wallis: F(2,15) = 10.3, *p* < 0.01; post hoc Steel-Dwass test, buffer vs Aβ42, *p* = 0.0170, at 19:00, Kruskal–Wallis: F(2,15) = 8.3, *p* < 0.05; post hoc Steel-Dwass test, Aβ40 vs Aβ42, *p* = 0.0202; Day9–11, at 22:00, Kruskal–Wallis: F(2,15) = 7.4, *p* < 0.05; post hoc Steel-Dwass test, buffer vs Aβ42, *p* = 0.0464. NREM: Day6–8, at 4:00, Kruskal–Wallis: F(2,15) = 11.4, *p* < 0.01; post hoc Steel-Dwass test, buffer vs Aβ40, *p* = 0.0402, buffer vs Aβ42, *p* = 0.0170, at 19:00, Kruskal–Wallis: F(2,15) = 7.8, *p* < 0.05; post hoc Steel-Dwass test, Aβ40 vs Aβ42, *p* = 0.0202; Day9–11, at 22:00, Kruskal–Wallis: F(2,15) = 7.1, *p* < 0.05; post hoc Steel-Dwass test, buffer vs Aβ42, *p* = 0.0464. REM: Day3–5, at 14:00, Kruskal–Wallis: F(2,15) = 7.1, *p* < 0.05; post hoc Steel-Dwass test, buffer vs Aβ42, *p* = 0.0286; Day6–8, at 8:00, Kruskal–Wallis: F(2,15) = 6.9, *p* < 0.05; post hoc Steel-Dwass test, Aβ40 vs Aβ42, *p* = 0.0310, at 9:00, Kruskal–Wallis: F(2,15) = 7.7, *p* < 0.05; post hoc Steel-Dwass test, buffer vs Aβ42, *p* = 0.0360; Day9–11, at 9:00, Kruskal–Wallis: F(2,15) = 6.3, *p* < 0.05; post hoc Steel-Dwass test, buffer vs Aβ42, *p* = 0.0281, at 12:00, Kruskal–Wallis: F(2,15) = 8.5, *p* < 0.05; post hoc Steel-Dwass test, buffer vs Aβ42, *p* = 0.0457, Aβ40 vs Aβ42, *p* = 0.0196. The gray background indicates the DP. Values are presented as means ± SEM; *, *p* < 0.05.

**Extended Figure 3. Alterations of sleep/wakefulness states 3 days to 11 days after Aβ injection**

(a–c) Total amount during 24 hours (a), during the DP (b), and during the LP (c) of wakefulness (left), NREM sleep (middle), and REM sleep (right), 3 days to 11 days after Aβ injection. REM: Day3–5: 24 hours, Kruskal–Wallis: F(2,15) = 7.2, *p* < 0.05; post hoc Steel-Dwass test, buffer vs Aβ42, *p* = 0.0170; LP, Kruskal–Wallis: F(2,15) = 6.7, *p* < 0.05; post hoc Steel-Dwass test, buffer vs Aβ42, *p* = 0.0170; Day6–8: LP, Kruskal–Wallis: F(2,15) = 6.1, *p* < 0.05; post hoc Steel-Dwass test, buffer vs Aβ42, *p* = 0.0464. (d–f) Episode number during 24 hours (d), during the DL (e), and during the LP (f) of wakefulness (left), NREM sleep (middle), and REM sleep (right), 3 days to 11 days after Aβ injection. REM: Day3–5: 24 hours, Kruskal–Wallis: F(2,15) = 6.1, *p* < 0.05; post hoc Steel-Dwass test, buffer vs Aβ42, *p* = 0.0170; LP, Kruskal–Wallis: F(2,15) = 6.3, *p* < 0.05; post hoc Steel-Dwass test, buffer vs Aβ42, *p* = 0.0170. (g–i) Mean episode duration during 24 hours (g), during the DP (h), and during the LP (i) of wakefulness (left), NREM sleep (middle), and REM sleep (right), 3 days to 11 days after Aβ injection. The gray background indicates the DP. The yellow background indicates the LP. Values are presented as means ± SEM; *, *p* < 0.05.

**Extended Figure 4. Frequency of sleep/wakefulness stage transitions 3 days to 11 days after Aβ injection**

(a–f) Frequency of sleep/wakefulness stage transitions during 24 hours after Aβ injection. The number of transitions between all states during 24 hours for 3 to 5 days (a), 6 to 8 days (c), and 9 to 11 days (e) following Aβ administration. The number of transitions between each sleep/wakefulness state for 3 to 5 days (b), 6 to 8 days (d), and 9 to 11 days (f) following Aβ administration. Left, buffer-treated mice; middle, Aβ40-treated mice; right, Aβ42-treated mice. Day3–5: from NREM to REM: Kruskal–Wallis: F(2,15) = 6.1, *p* < 0.05; post hoc Steel-Dwass test, buffer vs Aβ42, *p* = 0.0170. (g–l) Frequency of sleep/wakefulness sage transitions during the DP after Aβ injection. The number of transitions between all states during the dark period for 3 to 5 days (g), 6 to 8 days (i), and 9 to 11 days (k) following Aβ administration. The number of transitions between each sleep/wakefulness state for 3 to 5 days (h), 6 to 8 days (j), and 9 to 11 days (l) following Aβ administration. Left, buffer-treated mice; middle, Aβ40-treated mice; right, Aβ42-treated mice. (m–r) The frequency of sleep/wakefulness stage transitions during the LP after Aβ injection. The number of transitions between all states during the LP for 3 to 5 days (m), 6 to 8 days (o), and 9 to 11 days (q) following Aβ administration. The number of transitions between each sleep/wakefulness state for 3 to 5 days (n), 6 to 8 days (p), and 9 to 11 days (r) following Aβ administration. Left, buffer-treated mice; middle, Aβ40-treated mice; right, Aβ42-treated mice. Day3–5: From NREM to REM: Kruskal–Wallis: F(2,15) = 6.3, *p* < 0.05; post hoc Steel-Dwass test, buffer vs Aβ42, *p* = 0.0170. Day9–11: From NREM to wakefulness: Kruskal–Wallis: F(2,15) = 6.1, *p* < 0.05; post hoc Steel-Dwass test, buffer vs Aβ42, *p* = 0.0464. NR, NREM sleep; R, REM sleep; W, wakefulness. The gray background indicates the DP. The yellow background indicates the LP. Values are presented as means ± SEM; *, *p* < 0.05.

**Extended Figure 5. EEG power spectra of sleep/wakefulness states during over a 24-hour period, from 3 to 11 days after Aβ injection**

(a, e, and i) Average EEG power spectra during wakefulness (a), NREM sleep (e), and REM sleep (i) for 3 to 5 days (left), 6 to 8 days (middle), and 9 to 11 days (right) following Aβ administration. (b–d, f–g, and j–l) Comparison of normalized delta (1–5 Hz) power (b, f, and j), theta (6–10 Hz) power (c, g, and k), and ratio of theta to delta power (d, h, and l) during wakefulness (b–d), NREM sleep (f–h), and REM sleep (j–l). Wakefulness: delta: Day 6–8: Kruskal–Wallis: F(2,14) = 7.1, *p* < 0.05; post hoc Steel-Dwass test, buffer vs Aβ40, *p* = 0.0155, Day 9–11: Kruskal–Wallis: F(2,14) = 11.4, *p* < 0.01; post hoc Steel-Dwass test, buffer vs Aβ40, *p* = 0.0300, Aβ40 vs Aβ42, *p* = 0.0152: theta: Day 3–5: Kruskal–Wallis: F(2,14) = 7.0, *p* < 0.05; post hoc Steel-Dwass test, buffer vs Aβ42, *p* = 0.0458, Day 6–8: Kruskal–Wallis: F(2,14) = 10.4, *p* < 0.01; post hoc Steel-Dwass test, buffer vs Aβ40, *p* = 0.0155, buffer vs Aβ42, *p* = 0.0240, Day 9–11: Kruskal–Wallis: F(2,14) = 8.9, *p* < 0.05; post hoc Steel-Dwass test, buffer vs Aβ40, *p* = 0.0188, Aβ40 vs Aβ42, *p* = 0.0294: theta/delta ratio: Day 6–8: Kruskal–Wallis: F(2,14) = 10.0, *p* < 0.01; post hoc Steel-Dwass test, buffer vs Aβ40, *p* = 0.0123, buffer vs Aβ42, *p* = 0.0240, Day 9–11: Kruskal–Wallis: F(2,14) = 10.6, *p* < 0.01; post hoc Steel-Dwass test, buffer vs Aβ40, *p* = 0.0188. REM: delta: Day 9–11: Kruskal–Wallis: F(2,14) = 7.0, *p* < 0.05; post hoc Steel-Dwass test, buffer vs Aβ42, *p* = 0.0313: theta: Day 3–5: Kruskal–Wallis: F(2,14) = 6.9, *p* < 0.05; post hoc Steel-Dwass test, buffer vs Aβ42, *p* = 0.0245, Day 6–8: Kruskal–Wallis: F(2,14) = 7.9, *p* < 0.01; post hoc Steel-Dwass test, buffer vs Aβ42, *p* = 0.0240, Day 9–11: Kruskal–Wallis: F(2,14) = 9.4, *p* < 0.01; post hoc Steel-Dwass test, buffer vs Aβ40, *p* = 0.0188, buffer vs Aβ42, *p* = 0.0218: theta/delta ratio: Day 9–11: Kruskal–Wallis: F(2,14) = 8.7, *p* < 0.05; post hoc Steel-Dwass test, buffer vs Aβ42, *p* = 0.0313. Values are presented as means ± SEM; *, *p* < 0.05.

**Extended Figure 6. EEG power spectra of sleep/wakefulness states during the DP, from 3 to 11 days after Aβ injection**

(a, e, and i) Average EEG power spectra during wakefulness (a), NREM sleep (e), and REM sleep (i) for 3 to 5 days (left), 6 to 8 days (middle), and 9 to 11 days (right) following Aβ administration. (b–d, f–g, and j–l) Comparison of normalized delta (1–5 Hz) power (b, f, and j), theta (6–10 Hz) power (c, g, and k), and ratio of theta to delta power (d, h, and l) during wakefulness (b–d), NREM sleep (f–h), and REM sleep (j–l). Wakefulness: delta: Day 6–8: Kruskal–Wallis: F(2,14) = 7.3, *p* < 0.05; post hoc Steel-Dwass test, buffer vs Aβ40, *p* = 0.0144, Day 9–11: Kruskal–Wallis: F(2,14) = 9.3, *p* < 0.01; post hoc Steel-Dwass test, buffer vs Aβ40, *p* = 0.0181, Aβ40 vs Aβ42, *p* = 0.0188: theta: Day 6–8: Kruskal–Wallis: F(2,14) = 9.8, *p* < 0.01; post hoc Steel-Dwass test, buffer vs Aβ40, *p* = 0.0188, buffer vs Aβ42, *p* = 0.0240, Day 9–11: Kruskal–Wallis: F(2,14) = 6.4, *p* < 0.05; post hoc Steel-Dwass test, buffer vs Aβ40, *p* = 0.0456: theta/delta ratio: Day 6–8: Kruskal–Wallis: F(2,14) = 10.3, *p* < 0.01; post hoc Steel-Dwass test, buffer vs Aβ40, *p* = 0.0125, buffer vs Aβ42, *p* = 0.0245, Day 9–11: Kruskal–Wallis: F(2,14) = 8.1, *p* < 0.05; post hoc Steel-Dwass test, buffer vs Aβ40, *p* = 0.0247. REM: delta: Day 9–11: Kruskal–Wallis: F(2,14) = 6.4, *p* < 0.05; post hoc Steel-Dwass test, buffer vs Aβ42, *p* = 0.0422: theta: Day 3–5: Kruskal–Wallis: F(2,14) = 7.3, *p* < 0.05; post hoc Steel-Dwass test, buffer vs Aβ42, *p* = 0.0240, Day 6–8: Kruskal–Wallis: F(2,14) = 6.5, *p* < 0.05; post hoc Steel-Dwass test, buffer vs Aβ42, *p* = 0.0414, Day 9–11: Kruskal–Wallis: F(2,14) = 9.6, *p* < 0.01; post hoc Steel-Dwass test, buffer vs Aβ40, *p* = 0.0306, buffer vs Aβ42, *p* = 0.034: theta/delta ratio: Day 9–11: Kruskal–Wallis: F(2,14) = 8.7, *p* < 0.05; post hoc Steel-Dwass test, buffer vs Aβ40, *p* = 0.0313, buffer vs Aβ42, *p* = 0.0234. Values are presented as means ± SEM; *, *p* < 0.05.

**Extended Figure 7. EEG power spectra of sleep/wakefulness states during the LP, from 3 to 11 days after Aβ injection**

(a, e, and i) Average EEG power spectra during wakefulness (a), NREM sleep (e), and REM sleep (i) during days 3 to 5 (left), 6 to 8 (middle), and 9 to 11 after Aβ injection. (b–d, f–g, and j–l) Comparison of normalized delta (1–5 Hz) power (b, f, and j), theta (6–10 Hz) power (c, g, and k), and ratio of theta to delta power (d, h, and l) during wakefulness (b–d), NREM sleep (f–h), and REM sleep (j–l). Wakefulness: delta: Day 6–8: Kruskal–Wallis: F(2,14) = 6.5, *p* < 0.05; post hoc Steel-Dwass test, buffer vs Aβ40, *p* = 0.0157, Day 9–11: Kruskal–Wallis: F(2,14) = 12.0, *p* < 0.01; post hoc Steel-Dwass test, buffer vs Aβ40, *p* = 0.0200, Aβ40 vs Aβ42, *p* = 0.0123: theta: Day 6–8: Kruskal–Wallis: F(2,14) = 10.2, *p* < 0.01; post hoc Steel-Dwass test, buffer vs Aβ40, *p* = 0.0117, buffer vs Aβ42, *p* = 0.0234, Day 9–11: Kruskal–Wallis: F(2,14) = 9.1, *p* < 0.05; post hoc Steel-Dwass test, buffer vs Aβ40, *p* = 0.0181, buffer vs Aβ42, *p* = 0.0287: theta/delta ratio: Day 6–8: Kruskal–Wallis: F(2,14) = 9.6, *p* < 0.01; post hoc Steel-Dwass test, buffer vs Aβ40, *p* = 0.0125, buffer vs Aβ42, *p* = 0.0320, Day 9–11: Kruskal–Wallis: F(2,14) = 11.0, *p* < 0.01; post hoc Steel-Dwass test, buffer vs Aβ40, *p* = 0.0202, Aβ40 vs Aβ42, *p* = 0.0390. REM: theta: Day 6–8: Kruskal–Wallis: F(2,14) = 8.0, *p* < 0.05; post hoc Steel-Dwass test, buffer vs Aβ42, *p* = 0.0240, Day 9–11: Kruskal–Wallis: F(2,14) = 9.5, *p* < 0.01; post hoc Steel-Dwass test, buffer vs Aβ40, *p* = 0.0183, buffer vs Aβ42, *p* = 0.0234: theta/delta ratio: Day 6–8: Kruskal–Wallis: F(2,14) = 7.7, *p* < 0.05; post hoc Steel-Dwass test, buffer vs Aβ42, *p* = 0.0245, Day 9–11: Kruskal–Wallis: F(2,14) = 8.7, *p* < 0.05; post hoc Steel-Dwass test, buffer vs Aβ40, *p* = 0.0314, buffer vs Aβ42, *p* = 0.0245. Values are presented as means ± SEM; *, *p* < 0.05.

**Extended Movie 1. Rapid eye movement during REM sleep of theta power-decreasing mice**

The video shows rapid eye movement during REM sleep in Aβ40-treated mice in the head-fixed condition. The box in the video (top right) shows the EEG (upper trace), EMG (lower trace), and power spectrum of every epoch. Blue, green, yellow, and orange bars represent delta, theta, alpha, and beta power, respectively. S, NREM sleep; R, REM sleep; W, wakefulness

