## Supplementary figures and images for "Amyloid fibrils in Alzheimer’s disease differently modulate sleep and cortical oscillations in mice depending on the type of amyloid"

### Supplementary Figure 1

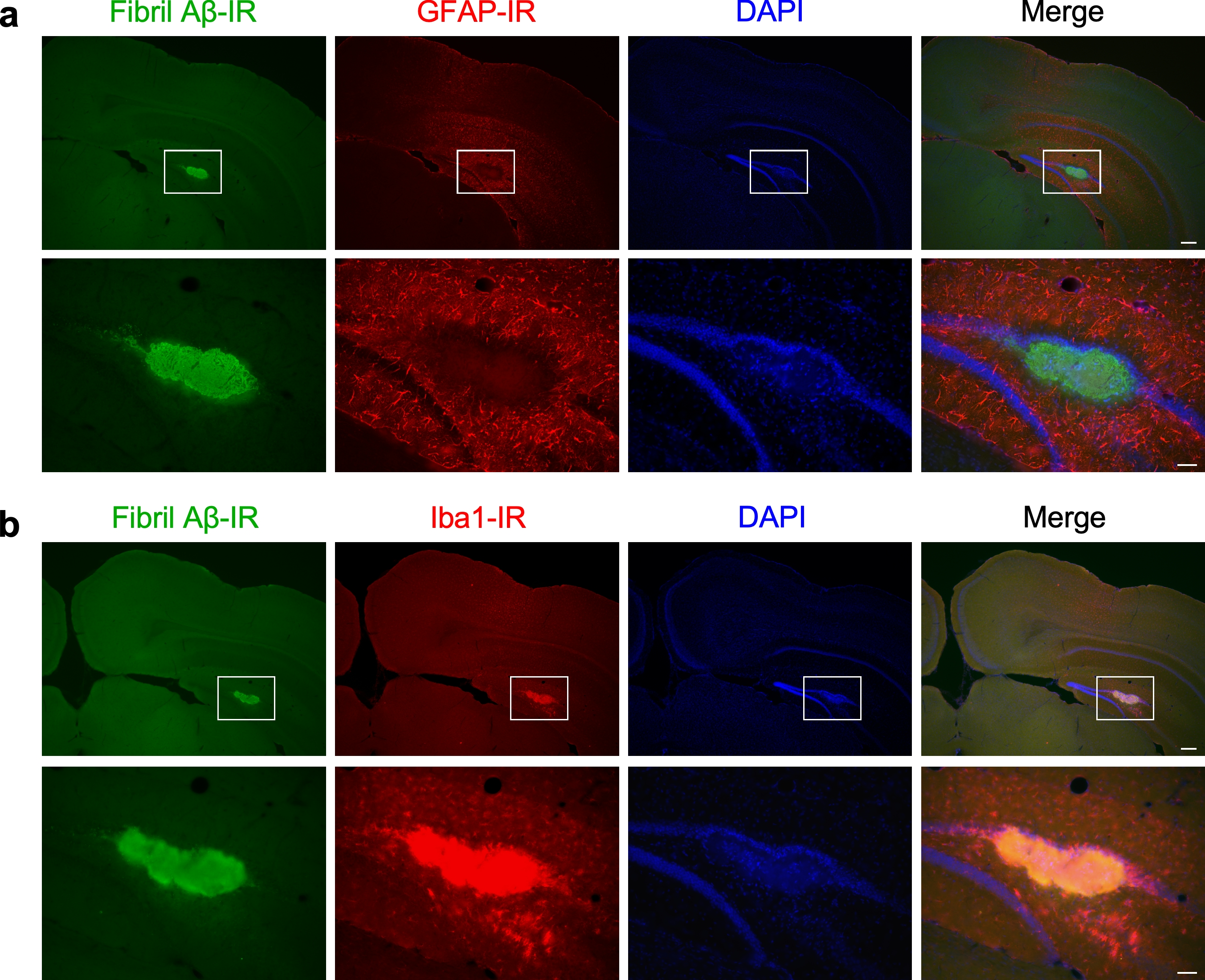

### Supplementary Figure 2

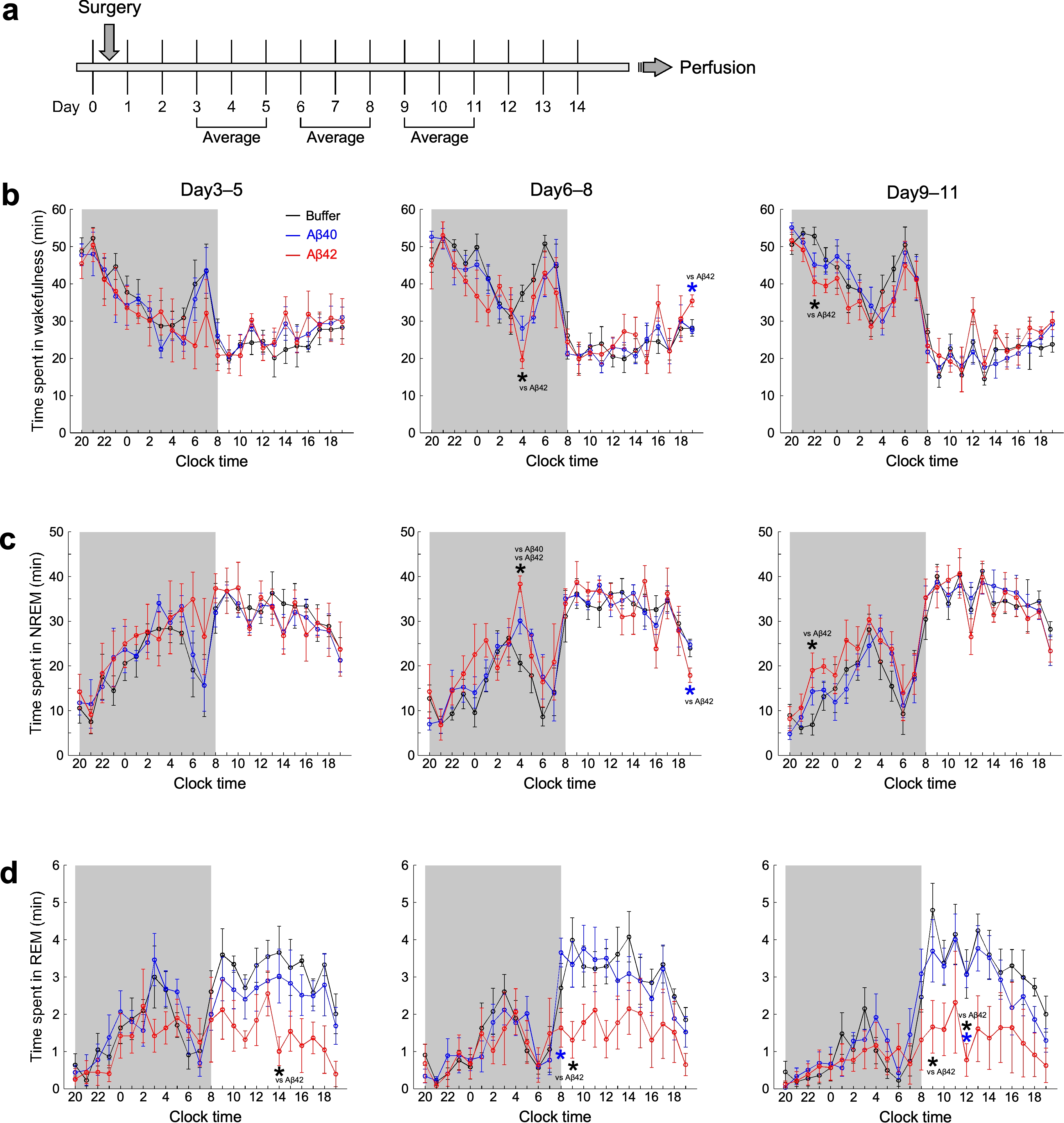

### Supplementary Figure 3

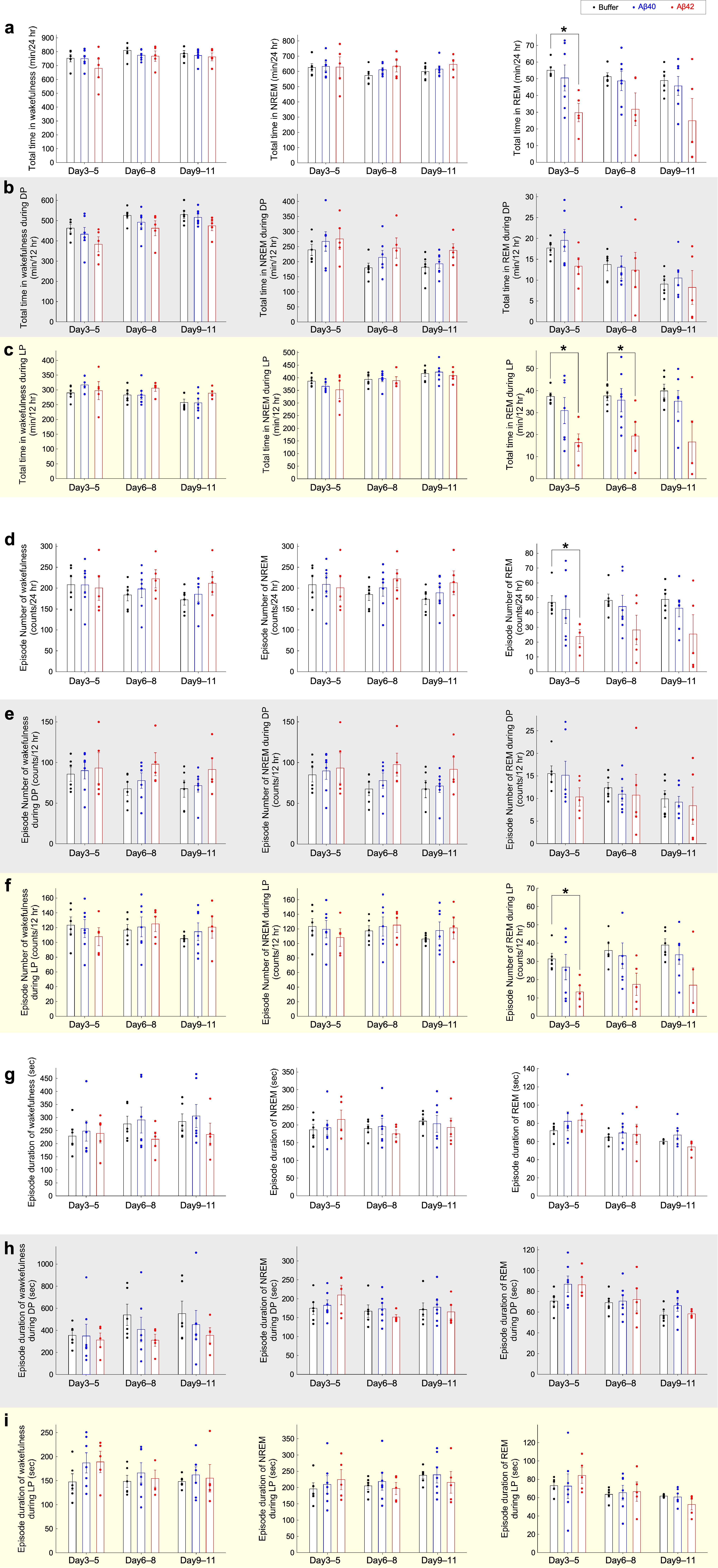

### Supplementary Figure 4

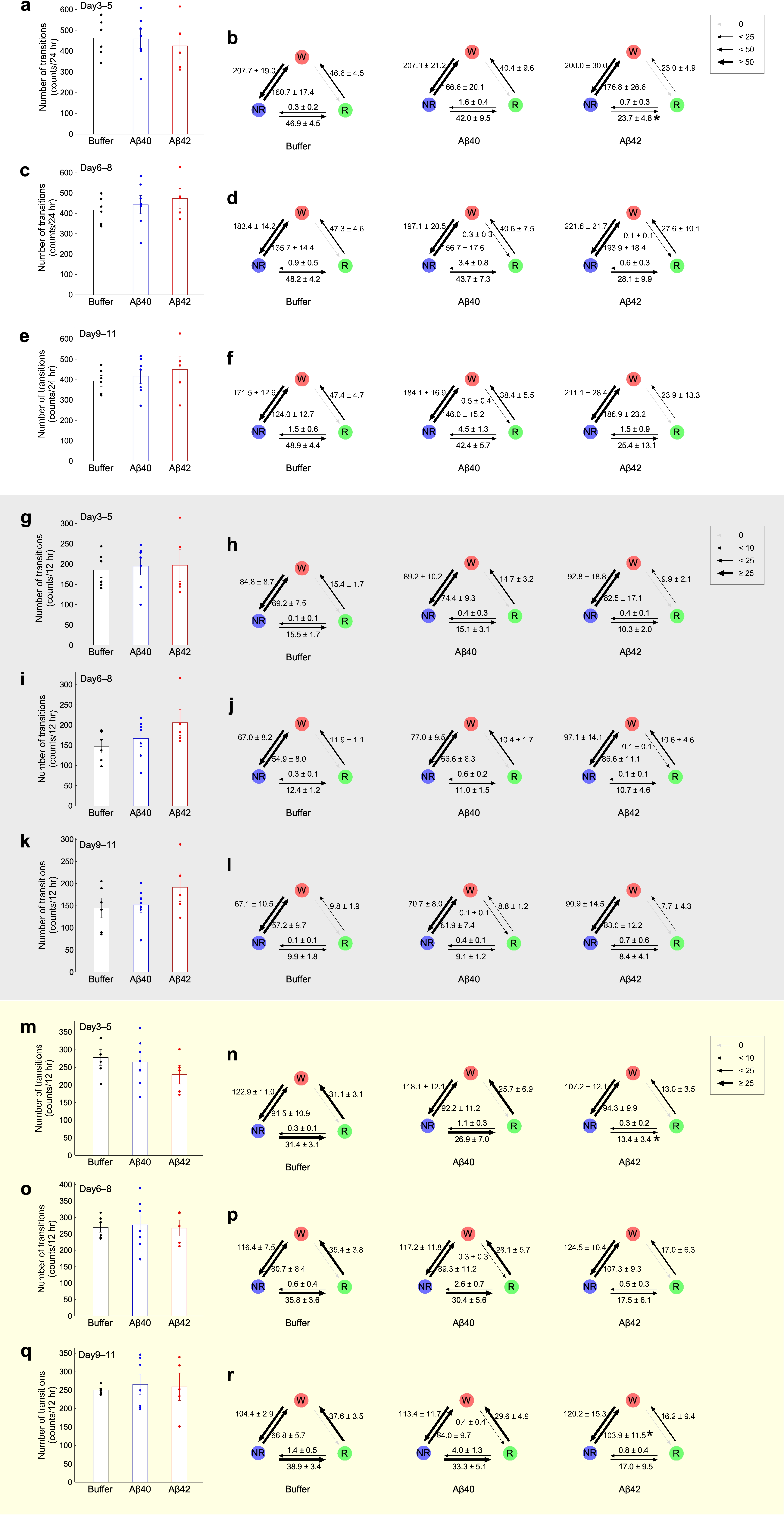

### Supplementary Figure 5

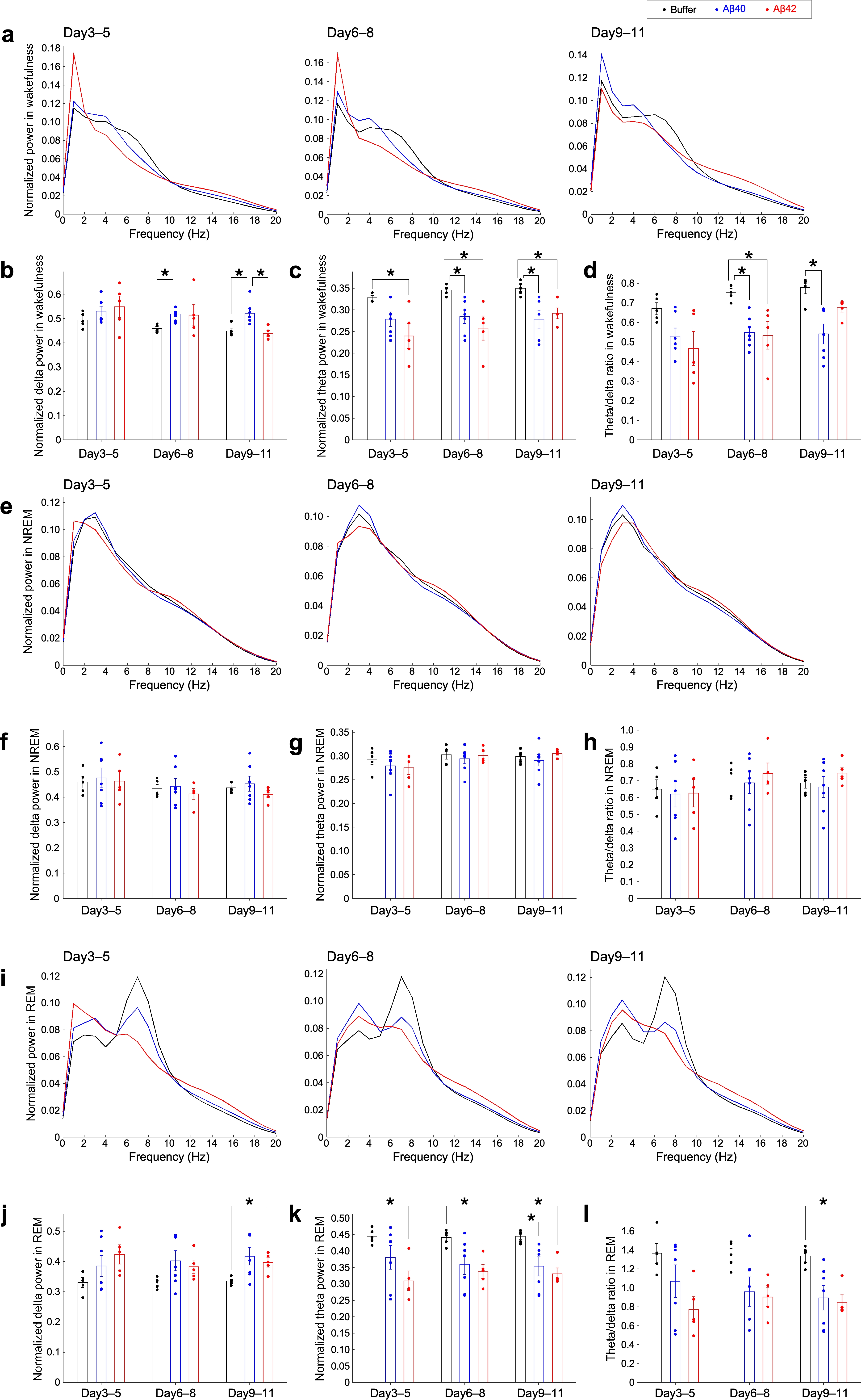

### Supplementary Figure 6

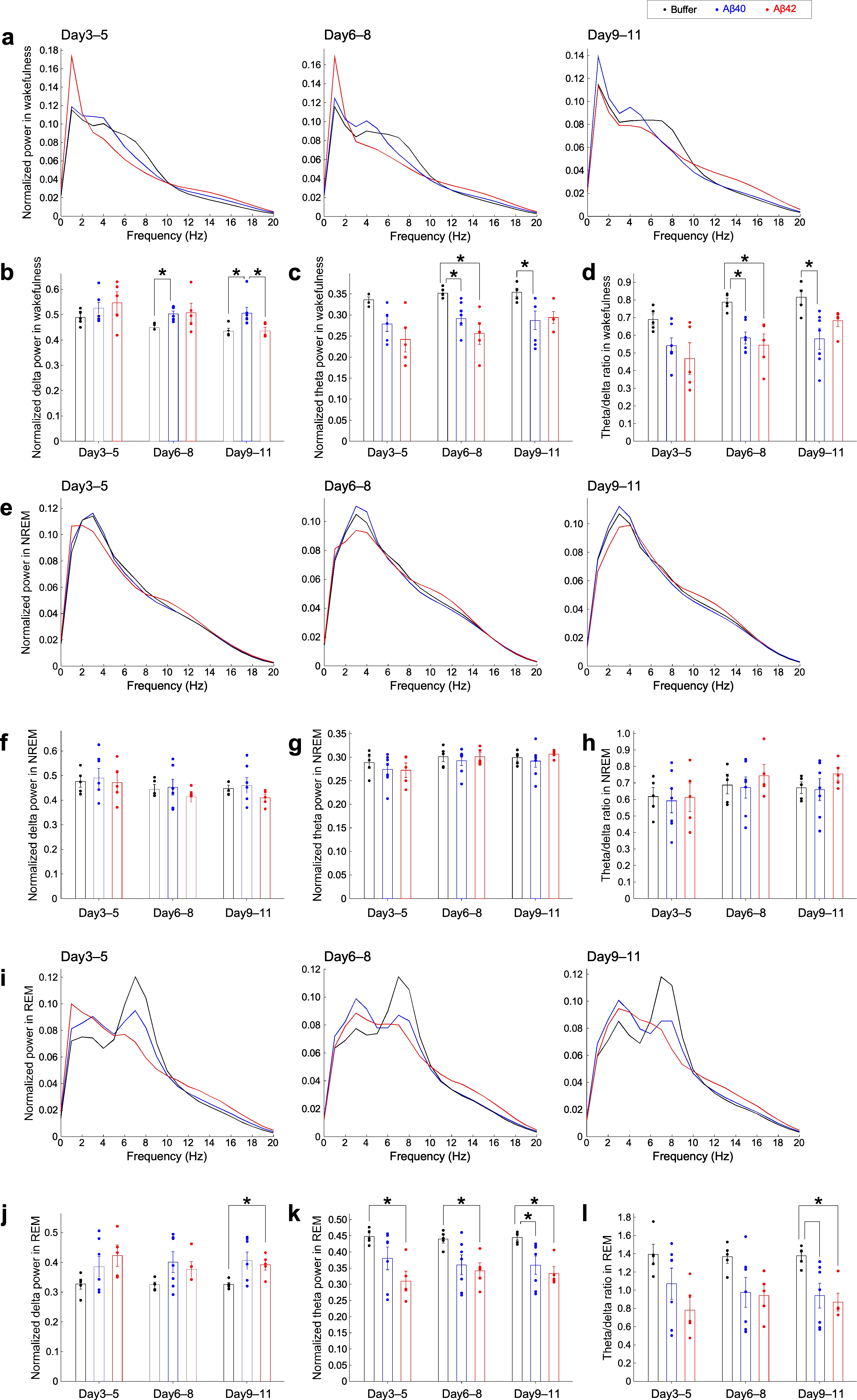

### Supplementary Figure 7

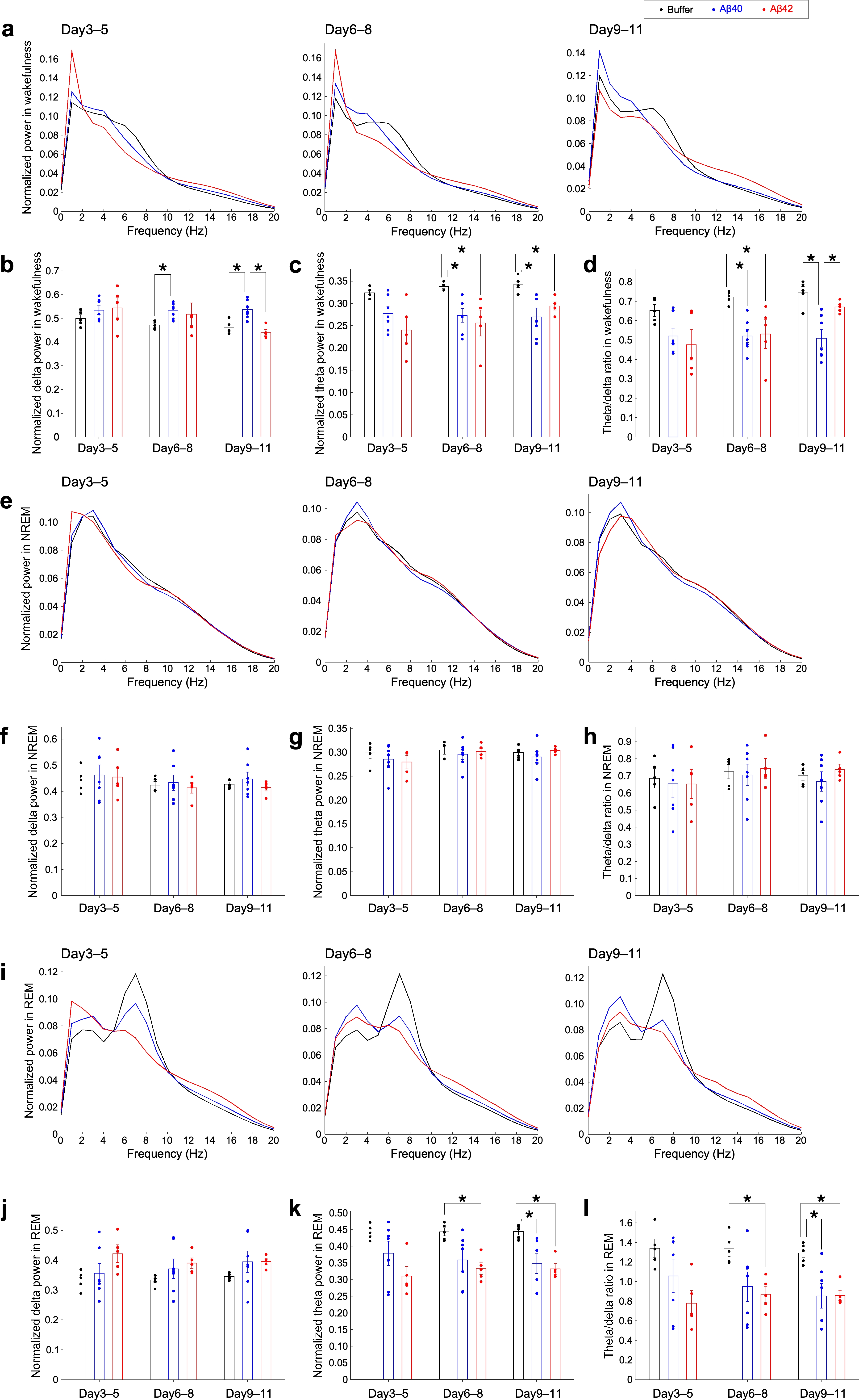
