## Supplementary material for "Amyloid fibrils in Alzheimer’s disease differently modulate sleep and cortical oscillations in mice depending on the type of amyloid": Methods

**Thioflavin T fluorescence assay**

The kinetic profiles of amyloid-beta1–40 (Aβ40) and amyloid-beta1–42 (Aβ42) amyloid formation were monitored using thioflavin T (ThT). Aβ peptides at 20 and 100 μM, prepared as described in our previous studies^1,2^, were incubated with 5 μM ThT at 37 °C in 20 mM 4-(2-hydroxyethyl)-1-piperazineethanesulfonic acid (HEPES) (pH 7.5) containing 100 mM NaCl. The samples were plated in triplicate onto a 96-well microplate with a nonbinding surface (Corning, product number: 3993). ThT fluorescence was measured every 5 min for 4 h using a SpectraMax iD3 Microplate Reader (Molecular Devices). The excitation and emission wavelengths were 445 nm and 490 nm, respectively. Shaking at medium strength for 250 s was applied between each reading.

ThT fluorescence data were fitted to the following equation to determine the lag time and elongation rate constant;

$$Y=y_{i}\text{ +}{\text{ }\text{m}}_{\text{i}}\text{t}\text{ + }\frac{\text{y}_{\text{f}}\text{ + }\text{m}_{\text{f}}\text{t}}{\text{1 + exp}\left[ \text{-}\text{k}\left( \text{t}\text{ - }\text{t}_{\text{0}} \right) \right]}$$

In which *Y* represents ThT fluorescence as a function of reaction time (*t*), *t*_0_ is the time when ThT fluorescence reaches half of its maximum value,*y*_i_ + *m*_i_*t* and *y*_f_ + *m*_f_*t* denote the initial and final baselines, respectively, *k* is the rate constant of elongation, and *t_0_* is the half time. The lag time is calculated using the relationship, *t*_lag_ = *t*_0_ - 2(1/*k*).

**Fluorescence dye-binding assay**

Stock solutions of ThT at 500 μM, 8-anilino-1-naphthalenesulfonic acid (ANS) at 500 μM, and curcumin at 2 mM were prepared using double-distilled water (ddH_2_O), ddH_2_O, and dimethyl sulfoxide (DMSO), respectively. Working solutions for the three dyes were freshly prepared by diluting the stocks in a buffer containing 20 mM HEPES (pH 7.5) and 100 mM NaCl. Prior to fluorescence measurements, 30 μL of Aβ40 or Aβ42 amyloid fibril solution was mixed with 70 μL of the above working solutions. The final concentration of both Aβ40 and Aβ42 was 30 μM, and the final concentrations of ThT, ANS, and curcumin were 10, 25, and 20 μM, respectively. The excitation wavelengths used were 445 nm for ThT, 370 nm for ANS, and 440 nm for curcumin. Fluorescence emission spectra were obtained at 37 °C using a Cary Eclipse fluorescence spectrophotometer (Agilent Technologies Inc.).

**Tyrosine fluorescence spectroscopy**

Amyloid samples were prepared using 40 μM of Aβ40 and Aβ42 in 20 mM HEPES buffer (pH 7.5) containing 100 mM NaCl. Tyrosine fluorescence spectra recorded from 290 to 375 nm were obtained using a Cary Eclipse fluorescence spectrophotometer (Agilent Technologies Inc.) with excitation at 275 nm. The slit widths of excitation and emission were 5 and 10 nm, respectively. The path length of the quartz cuvette was 1 cm. All measurements were performed at 37 °C.

**One-dimensional proton NMR spectroscopy**

One-dimensional (1D) proton (^1^H) NMR spectra of Aβ40 and Aβ42 samples were obtained using a Bruker Avance II 800 NMR spectrometer equipped with a cryogenic probe (Bruker BioSpin) at 10 °C. A standard pulse sequence with water suppression by excitation sculpting (zgesgp) from Bruker Topspin was used. Aβ40 and Aβ42 samples at 25 μM were prepared in 20 mM HEPES buffer (pH 7.5) containing 100 mM NaCl and 10% Deuterium oxide (D_2_O) (*v*/*v*). Each spectrum was obtained from 512 transients. Data were processed using TopSpin 3.6.1 software (Bruker Topspin).

**Atomic force microscopy**

Sample solutions (20 μL) were applied to a freshly cleaved mica plate, which was then incubated for 10 min. Each sample was gently rinsed twice with 100 μL of distilled water. The remaining water was carefully removed using filter paper and compressed air. Measurements and analyses of atomic force microscopy (AFM) were conducted using Park NX10 (Park Systems Corp.). Data were analyzed using Gwyddion software^3^.

**Transmission electron microscopy**

Transmission electron microscopy (TEM) images were obtained using a Bio-High voltage EM system (JEM-1400 Plus at 120 kV, JEOL Ltd.). A sample solution (5 μL) was applied to a collodion-coated copper grid (Nisshin EM Co.), and incubated it for 1 min. The remaining solution was then removed with filter paper, and 5 μL of distilled water containing 2% (*w*/*w*) uranyl acetate was spotted onto the grid. After 1 min, the remaining solution was removed in the same manner. Data were analyzed using ImageJ software^4^.

**Circular dichroism spectroscopy**

Far-UV circular dichroism (CD) measurements of Aβ40 and Aβ42 amyloid fibrils were performed on a JASCO J-710 spectrophotometer (JASCO Corporation) using a quartz cuvette with a 1 mm path length at room temperature. Aβ40 or Aβ42 at 20 μM was dissolved in 20 mM HEPES buffer (pH 7.5) containing 100 mM NaCl, and incubated for amyloid formation. The bandwidth and scan rate were 1 nm and 100 nm min^−1^, respectively. CD signals are presented as the mean residue ellipticity (deg cm^2^ dmol^−1^). The content of secondary structures was predicted using the method reported previously^5^.

**Mice**

All experimental procedures involving animals were approved by the Animal Care and Use Committee of Tohoku University (approval no.: 2021LsA-001), and Hokkaido University (approval no.: 23-0117) and were conducted in accordance with the National Institutes of Health guidelines. All efforts were made to minimize animal suffering and discomfort, and to reduce the number of animals used. Mice were housed under a controlled 12 h/12 h light/dark cycle (light on hours: 8:00­­–20:00). Mice had *ad libitum* access to food and water. A total of 21 male wild-type C57BL/6J mice (CLEA Japan) were used in this study. For freely moving EEG/EMG recordings, 18 mice were used. For head-fixed EEG/EMG/pupil recordings, two mice were used. For immunohistochemical analysis, 13 animals used. In this study, female mice were not used because the sleep pattern of mice on the C57 genetic background has been reported to be affected by the estrous cycle^6^.

**Surgical procedures**

Male wild-type C57BL/6J mice of about 3-months old were used (94.8 ± 3.2 days of age). Stereotaxic surgery was performed under anesthesia with pentobarbital (5 mg/kg, intraperitoneal injection as induction) and with isoflurane (1%–2% for maintenance) using a vaporizer for small animals (Bio Research Center) with the mice positioned in a stereotaxic frame (Narishige). The detailed procedure of intrahippocampal injection has been described in our previous paper^1^. Briefly, Aβ40, Aβ42, or buffer (20 mM HEPES buffer [pH 7.5], 100 mM NaCl) was bilaterally injected into the hippocampus (2.5 mm posterior, ± 2.0 mm lateral, and 2.0 mm depth from bregma) in a volume of 1.5 µL, at a flow rate of 0.2 µL/min using a Hamilton 10-µL syringe. The injection volume and flow rate were controlled using a stereotaxic injector (Legato 130, Muromachi Kikai). For EEG/EMG recordings, three bone screws were implanted into the skull as electrodes for cortical EEGs, and twisted wires (AS633, Cooner wire) were inserted into the neck muscle as electrodes for EMGs. Another bone screw was implanted into the cerebellum as a ground. All electrodes were connected to a pin socket and fixed to the skull with dental cement. For head-fixed recordings, a stainless chamber (CF-10, Narishige) was also attached to the skull.

***In vivo* sleep/wakefulness recordings using freely moving mice**

Continuous EEG and EMG recordings were conducted through a slip ring (SPM-35-8P-03, Hikari Denshi), which was designed so that the movement of the mice was unrestricted. Two of the three screw electrodes were used for EEG recordings. Differential amplification was performed for EEG and EMG signals (AB-610J, Nihon Koden). Amplified signals were then filtered (EEG, 0.75–20 Hz; EMG, 20–50 Hz), digitized at a sampling rate of 128 Hz, and recorded using SleepSign software version 3 (Kissei Comtec). Behavior of the mice was monitored through a charge coupled device (CCD) video camera, and recorded onto a computer synchronized with EEG and EMG recordings using the SleepSign video option system (Kissei Comtec).

**Pupil recordings**

For habituation of the head-fixed condition, mice were placed in a head-fix apparatus (MAG-1, Narishige) by securing them using a stainless chamber frame and placing them into an acrylic tube. This procedure was continued for at least 5 days, during which the duration of head-fixation was gradually extended from 10 to 120 min. To record rapid eye movement during REM sleep, mouse pupils were monitored with an off-axis infrared (IR) light source (940 nm IR light-emitting diode [LED]). A camera (BU130, Toshiba Teli) with a zoom lens (HF25HA-1B, Fujifilm) and an IR filter (R-72, Edmund) was placed approximately 10 cm from the mouse’s right eye. Images were collected at 30 Hz using custom-written MATLAB software (MathWorks).

**Immunohistochemistry**

To confirm the localization of the amyloid fibrils, microglia, and astrocytes, mice were deeply anesthetized with isoflurane, and perfused sequentially with 20 mL of chilled phosphate-buffered saline (PBS) and 20 mL of chilled 4% paraformaldehyde in PBS (Nacalai Tesque). Brains were removed and immersed in the above fixation solution overnight at 4 °C, and then immersed in 30% sucrose in PBS for at least 2 days. The brains were then quickly frozen in embedding solution (Sakura Finetek), and cut into coronal sections using a cryostat (CM1850, Leica) at a thickness of 40 µm. For immunostaining, the brain sections were incubated with an anti-amyloid fibril antibody (1:500; ab126468, Abcam) and anti-GFAP antibody (1:2,000; G3893, Sigma) overnight at 4 °C. Then, the sections were incubated with CF488A donkey anti-rabbit Immunoglobulin G (IgG) (1:1,000; 20015-1, Nacalai Tesque) and CF594 donkey anti-mouse IgG (1:1,000; 20115-1, Nacalai Tesque) for 1 h at room temperature. For microglia immunostaining, the brain sections were incubated with SPICA Dye568-conjugated anti-Iba1 antibody (1:200; 011-28013, Fujifilm Wako Pure Chemical Co.) overnight at 4 °C. The sections were incubated with DAPI (1:1,000; D523, Dojindo) for 30 min at room temperature, and then mounted onto aminosilane (APS) -coated slides, coverslipped with 50% glycerol in PBS, and observed using a fluorescence microscope (BZ-9000, Keyence).

**Immunohistochemical analysis**

To quantify Aβ-induced hippocampal neuronal loss, color intensity analysis was performed using ImageJ software. One in 6 of the series of brain sections from 1.82 mm to 2.7 mm posterior from bregma were selected. The entire hippocampus was selected as a region of interest (ROI). A light-intensity threshold was set, and the percentage of area above the threshold in the ROI was calculated. The average value was then calculated and quantified for each mouse.

**Sleep scoring and EEG analysis**

Polysomnographic recordings were automatically scored offline, with each epoch scored as wakefulness, NREM sleep, or REM sleep by SleepSign (Kissei Comtec), in 4-sec epochs, according to standard criteria^7,8^. All vigilance state classifications assigned by SleepSign were confirmed visually. The same individual, blinded to experimental condition, scored all EEG/EMG recordings. Three consecutive 4-sec epochs (12 sec) needed to be scored as a single state before being considered as a sleep or wake bout, and used for further analysis. Spectral analysis of the EEGs was performed by fast Fourier transform, which yielded a power spectral profile with a 1-Hz resolution divided into delta (1−5 Hz), theta (6−10 Hz), alpha (10−13 Hz), and beta (13−20 Hz) waves. The EEG spectral power was normalized in the following way based on our previous papers^9^. EEGs from mice were averaged in 1-Hz bins for each of wakefulness, NREM sleep, and REM sleep. Then, each of the averages in 1-Hz bins were divided by the total of the average values from 0.75 to 20 Hz for each of wakefulness, NREM sleep, and REM sleep. To calculate the EEG power of the delta and theta bands for each sleep/wakefulness state, the EEG values from 1 to 5 Hz and 6 to 10 Hz were summed, respectively. Each summed value was then divided by the total average value from 0.75 to 20 Hz for wakefulness, NREM sleep, and REM sleep.

**Statistical analysis**

Data are presented as the mean ± SEM, unless otherwise stated. In Figs 3 to 5, and Extended figs, statistical analyses were performed using MATLAB software. For multiple comparisons of sleep/wakefulness patterns and EEG power, the Kruskal-Wallis test was performed with the *post hoc* Steel-Dwass test. In Fig. 6, one-way analysis of variance (ANOVA) with the post hoc Fisher’s protected least significant difference test was used. A *p*-value of less than 0.05 was considered to indicate a statistically significant difference between groups.

**Acknowledgements**

This work was supported partially by grants from the Frontier Research Institute for Interdisciplinary Sciences (FRIS) Creative Interdisciplinary Collaboration Program, the Uehara Memorial Foundation, the Astellas Foundation for Research on Metabolic Disorders, Daiichi Sankyo Foundation of Life Science, and Fusion Oriented Research for disruptive Science and Technology (FOREST) from Japan Science and Technology Agency (grant no.: JPMJFR2047), to T.T.; by grants from FOREST (grant no.: JPMJFR201F), the Takeda Science Foundation, the Mochida Memorial Foundation for Medical and Pharmaceutical Research, the Naito Foundation, the Uehara Memorial Foundation, the Terumo Life Science Foundation, the Astellas Foundation for Research on Metabolic Disorders, The Asahi Glass Foundation, Mitsui Sumitomo Insurance Welfare Foundation, the Daiichi Sankyo Foundation of Life Science, Sumitomo Foundation, Ono Medical Research Foundation, and Nakatani Foundation, to M.O.; by grants from Korea Basic Science Institute (KBSI) (A439200, C539200, C523200, C512120, A412580, and A423310) and the National Research Foundation of Korea funded by the Korean government (grant no.: RS-2022-NR069719) to Y.-H.L.; and the Sejong Science Fellowship Grant (grant no.: RS-2024-00356469) to Y.H. Generous support from the FRIS Cooperative Research Environment (CoRE), which is a shared research environment at Tohoku University, is acknowledged. We thank Professor Keizo Takao, Dr. Shoko Hashimoto, Dr. Eiko Minakawa, and Professor Shuzo Sakata for helpful discussions, and Dr. Helena Akiko Popiel for English language editing of the manuscript.
